# Inferring Single-Molecule Chromatin Interactions via Online Convex Network Dictionary Learning

**DOI:** 10.1101/2022.07.28.501904

**Authors:** Jianhao Peng, Chao Pan, Hanbaek Lyu, Minji Kim, Albert Cheng, Olgica Milenkovic

## Abstract

**Motivation:** Genomes of multicellular systems are compartmentalized and dynamically folded within the three-dimensional (3D) confines of the nucleus in order to facilitate gene regulation. Among the 3D-genome mapping technologies currently in use, droplet-based, barcode-linked sequencing (ChIA-Drop) has the unique capability to capture complex multi-way chromatin interactions at the single-molecule level. ChIA-Drop data gives rise to higher-order interaction networks in which nodes represent genomic fragments while (hyper)edges capture observed physical contacts. The problem of interest is to use this data to create a “dictionary” of interaction patterns (subnetworks) that accurately describe all global chromatin structures and associate dictionary elements with cellular functions.

**Results:** To construct interpretable chromatin dictionaries, we introduce a new algorithm termed online convex network dictionary learning (online cvxNDL). Unlike classical dictionary learning for image or text processing, online cvxNDL uses special subgraph sampling methods and produces interpretable subnetwork representatives corresponding to “convex mixtures” of patterns observed in real data. To demonstrate the utility of the method, we perform an in-depth study of RNAPII-enriched ChIA-Drop data from *Drosophila Melanogaster* S2 cell lines. Our results are two-fold: First, we show that online cvxNDL allows for accurate reconstruction of the original interaction network data using only a collection of roughly 25 dictionary elements and their “representatives” directly observed in the data. Second, we identify collections of interaction patterns of chromatin elements shared by related processes on different chromosomes and those unique to certain chromosomes. This is accomplished through Gene Ontology (GO) enrichment analysis that allows us to associate dictionary element representatives with functional properties of their corresponding chromatin region and in the process, determine what we call the “span” and “density” of chromatin interaction patterns.

**Availability and Implementation:** The code and dataset are available at: https://github.com/jianhao2016/online_cvxNDL/

## 1 Introduction

Eukaryotic genomes represent complex 3D structures that are compartmentalized and dynamically folded and unfolded within the nucleus. This topological organization of the genome plays an important role in cellular processes and gene regulation by allowing distal regulatory elements to enhance or suppress the expression of target genes^1,2^, and it has been widely studied using traditional “bulk” sequencing data^3,4^.

3D genome mapping technologies are used to record how various genomic loci engage in short- and long-range interactions. They include traditional Hi-C^3^, Micro-C^5^, used for capturing genome-wide unbiased chromatin topology, ChIA-PET^6,7^ and its variants PLAC-seq and HiChIP^8,9^, used for extracting chromatin interactions mediated by a specific protein factor. These methods effectively map the 3D structure of chromatin interactions onto a 2D contact map of different loci of the chromosome. However, due to the proximity ligation step, the methods can detect only pairwise contacts and hence fail to capture potential simultaneous interactions involving two or more genomic loci. Moreover, these technologies operate on a population of millions of molecules, thereby providing information about population averages only. To overcome such limitations, recent works have focused on developing ligation-free single-cell or single-molecule methods such as GAM^10^, (sc-)SPRITE^11,12^, and ChIA-Drop^13^.

Similar to commercially available scRNA-seq platforms, ChIA-Drop^13^ adopts a droplet-based barcode-linked technique to reveal multiway chromatin interactions at a single molecule level. The chromatin complexes are first encapsulated into gel-bead droplets, each with a unique barcode, and then sequenced and mapped to the reference genome. ChIA-Drop can simultaneously capture chromatin interactions of multiple loci and reveal the kinetics of loop formation^14^. ChIA-Drop data also reveals network patterns within single topologically associated domains (TADs) as well as some long-distance interactions across TADs (including ~ 15% of all chromatin complexes), which cannot easily be inferred from other data^13^. Single-molecule chromatin interactions can elucidate many cellular regulation and developmental phenomena. Still, at this point, *efficient* and *biologically interpretable* computational methods for analyzing long-distance multiplexed chromatin interactions at a single-cell or single-molecule level are lacking. Importantly, no associations between specific distal and proximal interactions topologies and cell functions are known.

Dictionary learning (DL), a form of (nonnegative) matrix factorization (MF), refers to learning a set of atoms (dictionary elements) that can approximate a matrix via (sparse) linear combinations of dictionary elements. DL is used for clustering, denoising and extraction of low-dimensional patterns from complex high-dimensional inputs^15–22^. For example, in image processing, dictionaries comprise collections of pixels whose linear combinations can be used to represent image patches (subimages). Standard DL methods^23,24^ have interpretability and scalability issues, and are mostly used with unstructured data. To address the interpretability issue, convex MF (CMF) was introduced in^25^. CMF requires the dictionary elements to be convex combinations of real data points. As an example, in the CMF setting, a dictionary element cannot be an arbitrary point that lacks a biological meaning or does not have a well connected topology. Instead, it has to be of the form of a convex combination of a small set of real data points. To scale the methods, DL and convex DL were adapted to an online setting^26,27^. DL for network-structured data was introduced in^28^. Network DL (NDL) works with subnetwork samples that are generated via Monte Carlo Markov Chain (MCMC) *subnetwork sampling*^28–30^. The gist of NDL is to identify a small number of network dictionary elements that best explain network interactions of the whole, global network in an efficient and accurate manner. Current online NDL algorithms do not provide directly interpretable results for biological networks.

We propose online cvxNDL, which is a new NDL method coupled with an MCMC sampling technique^29^ that also imposes “convexity constraints” on the sampled subnetworks and uses the notion of “dictionary element representatives”. The convex constraints force each learned dictionary element to be explainable through convex combination of a small subset of *real data subnetwork adjacency matrices.* For example, a dictionary element could be given as 50% of one observed interaction, and 50% of another. The two subset interactions constitute the *representatives* for the dictionary element. Hence, in the context of chromatin interaction networks, representatives are real data interaction subnetworks, and this allows one to use *Gene Ontology* (GO) enrichment analysis to uncover the joint functionality of genomic regions covering the representatives. Since GO terms are ordered hierarchically in the form of directed acyclic graphs^31^, with more general terms at the higher level closer to the root and more specific terms at the lower levels close to the leaves, the hierarchy can be used to select the most relevant (highest convex weight) representatives. These representatives, corresponding to real interaction patterns, can subsequently be associated with cellular functions. They may also be used to determine what we term “the span of interaction” (the largest linear genomic distance between interacting chromatin fragments) and “density of interaction” (which captures the density of interacting fragments within the span). Both concepts are rigorously defined in the Supplement Section 5.5.1.

We test our online cvxNDL algorithm on different chromosomes of the ChIA-Drop data of embryonic *Drosophila Melanogaster* Schneider 2 (S2) phagocytic cell lines, and provide biological interpretations of the chromatin interaction dictionary elements underlying certain developmental functions. Our results reveal different representative interaction patterns on L and R chromosomal arms, as well as different interaction spans and complexities for the 2R,L and 3R,L chromosomes. In addition to providing the first dictionaries of interaction in chromatin structures, online cvxNDL can be used in other areas of computational biology where the goal is to find small dictionaries of subnetwork interactions that describe a complex, global network. DL can also be used for compressing network data, which will be considered in a future work.

## 2 Methods

### Notation

Sets of consecutive integers are denoted by [*l*] = {1,..., *l*}. Capital letters are reserved for matrices (bold font) and random variables (RVs) (regular font). Vectors are denoted by lower-case underlined letters. For a matrix of dimension *d* × *n* over the reals, **A** ∈ ℝ^*d*×*n*^, **A**[*i*,:] is used to denote the *i*^th^ row and **A**[:, *i*] the *i*^th^ column of **A**. The entry in row *i*, column *j* is denoted by **A**[*i, j*]. Similarly, <u>*x*</u>[*l*] is used to denote the *l*^th^ coordinate of a deterministic vector *<u>x</u>* ∈ ℝ^*d*^. Furthermore, ||**A**||_1_ = **∑**_*i,j*_|**A**[*i, j*]| and 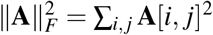.

A simple network 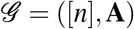 is an ordered pair of sets, the node set [*n*], and the set of edges given via their adjacency matrix **A**; here, **A**[*i, j*] = **A**[*j, i*] ∈ {0,1}, indicating the presence or absence of an undirected edge between vertices *i, j*. In addition, Col(**A**) stands for the set of columns of **A**, while cvx(**A**) stands for the convex hull of Col(**A**).

### Online DL

We first formulate the online DL problem. Assume that the input data samples are generated by a hidden random process and organized in matrices 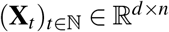 indexed by time *t*. For *n* = 1, **X**_*t*_ reduces to a column vector that encodes a *d*-dimensional signal. Given an (online, sequentially observed) data stream 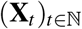, the goal is to find a sequence of dictionary matrices 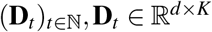, and codes 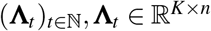, such that when *t* → ∞ almost surely we have

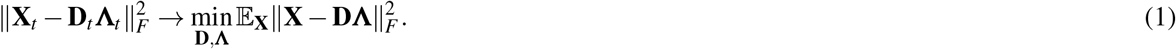

This expected loss in the previous equation can be minimized by iteratively by updating **Λ**_*t*_ and **D**_*t*_ every time a new data sample **X**_*t*_ is observed. The approximation error of **D** for a single data sample **X** and with sparsity-imposing regularizers is chosen as

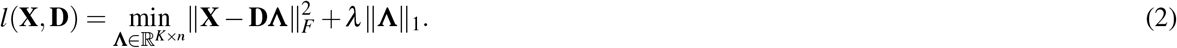

Furthermore, the empirical *f_t_* and surrogate loss 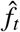 for **D** are defined as:

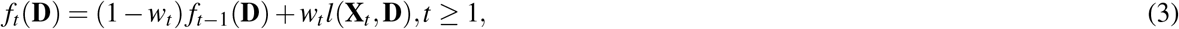

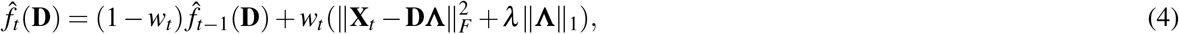

where *w_t_* is a weight that determines the sensitivity of the algorithm to the newly observed data. The online DL algorithm first updates the code matrix **Λ**_*t*_ by solving Equation (2) with *l*(**X**_*t*_, **D**_*t*–1_), then updates the dictionary matrix **D**_*t*_ by minimizing (4) via

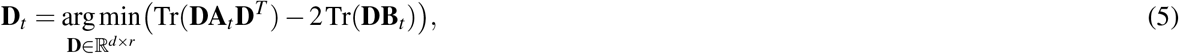

where 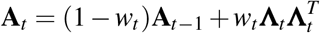 and 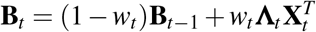 are the aggregated history of the input data and their codes. For simplicity, we set 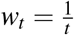.

To add convexity constraint to our dictionaries **D**_*t*_, we introduce for each dictionary element a representative set (region) 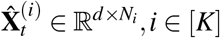, where *N_i_* is the size of the representative set for dictionary element **D**_*t*_ [:, *i*]. In a nutshell, the representative set for a dictionary element is a small subcollection of real data samples observed up to time t that best explain the dictionary element they are assigned to. The list of representative is updated after observing a sample the inclusion of which provides a better estimate of the dictionary element compared to the previous list. Since the representative list is bounded in size, if a new sample is included, an already existing sample has to be removed (see Figure 1). Formally, the optimization objective is of the form:

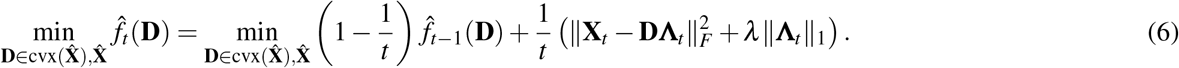

**Figure 1.**
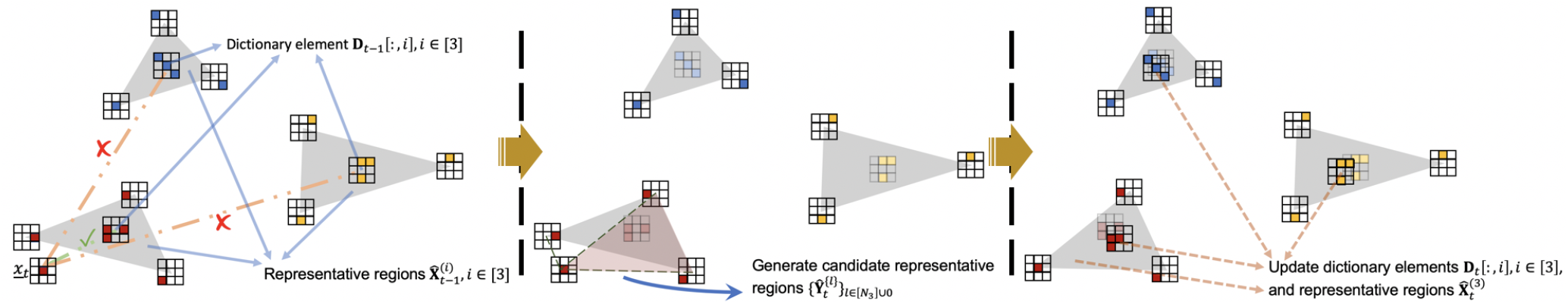
Illustration of the update procedure for the representative regions (lists) and dictionary elements. Upon observing a data sample online, the distance of the sample to each of the current dictionary elements is computed. Then, the sample is assigned to the representative region of the closest dictionary element. Data points in the corresponding representative region are updated to include or exclude the new sample according to the improvement or degradation in the quality of the dictionary element. Note that data points are adjacency matrices of real subnetworks.

### MCMC sampling of subnetworks

For NDL, it is natural to let the columns of **X**_*t*_ be vectorized adjacency matrices of *n* subnetworks. Hence it is required to efficiently sample meaningful subnetworks from large networks. For large networks, global random sampling is computationally demanding and produces mostly disconnected subnetworks. To address this problem, we proceed as follows. In image DL problems, samples can be generated directly from the image using adjacent rows and columns. However, such a sampling technique cannot be used for network data: Selecting nodes at random, along with their one-hop neighbors, may produce subnetworks of vastly different sizes and does not capture important long-range interactions. Also, it is difficult to determine how to trim these subnetworks. Sampling a fixed number of nodes uniformly at random from sparse networks and observing the induced subnetwork produces disconnected subnetworks with high probability. Instead, in this case, we consider “subnetwork sampling” introduced in^28,29^. Namely, we fix a template network *F* = ([*k*], **A**_*F*_) of *k* nodes, sample a random copy of *F* from 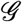 uniformly at random, and record the induced subnetwork. To achieve this, we use the MCMC sampling algorithm in^28^, which seeks subnetworks induced by *k* nodes in the original input network 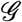, with the constraint that the subnetwork contains the template topology. Given an input network 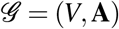 and a template network *F* = ([*k*], **A**_*F*_), we define a set of homomorphisms as a vector of the form (with the assumption that 0^0^ = 1):

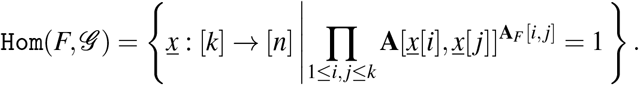

For each homomorphism 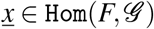, denote its induced adjacency matrix by **A**_<u>*x*</u>_, where **A**_<u>*x*</u>_[*a,b*] = **A**[<u>*x*</u>[*a*],<u>*x*</u>[*b*]], 1 ≤ *a, b* ≤ *k*. An example homomorphism is shown in Figure 2, where the input network 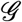 contains *n* = 9 nodes and the template network *F* is a star network that contains *k* = 4 nodes. One homomorphism in this case is <u>*x*</u>[*a*] = 9,*<u>x</u>*[*b*] = 6,<u>*x*</u>[*c*] = 4,*<u>x</u>*[*d*] = 7, which gives rise to an adjacency matrix **A**_<u>*x*</u>_ as depicted (for details, see the Supplement Section 5.2 Algorithm 1). Our choice of template network for subsequent analysis is a *k-chain*, a directed path from node 1 to *k*; chains are a simple and natural choice for networks with long average path lengths, such as chromatin interaction networks. This is the same choice of template as used in standard NDL.

**Figure 2.**
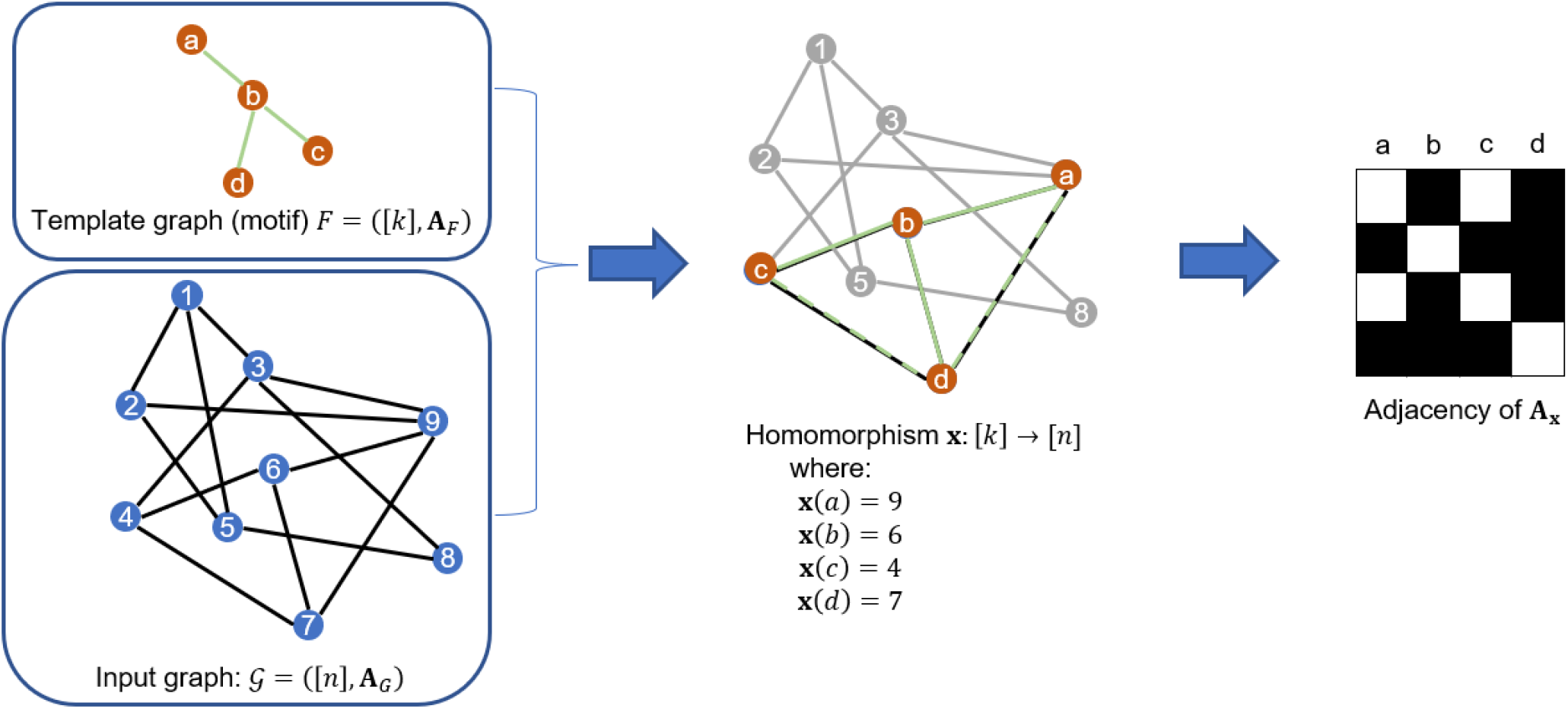
Subnetwork sampling and the notion of a homomorphism. In the adjacency matrix, a black field indicates 1, while a white field indicates 0.

**Algorithm 1.**
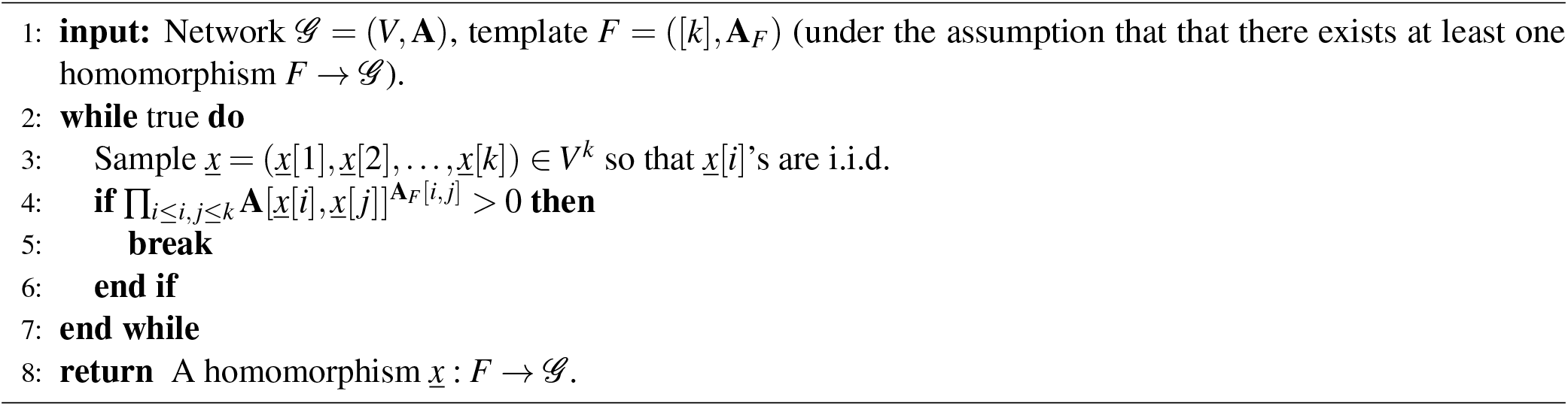
Rejection Sampling of Homomorphisms.

To efficiently generate a sequence of sample adjacency matrices 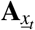 from 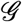 to use matrix samples **X**_*t*_, the MCMC sampling algorithm gradually changes the template network based on previous samples. An illustration of the sampling procedure is shown in Figure 3. In a nutshell, given a homomorphism 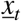 at step *t*, we first choose a node v from the neighborhood of 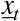 [1] with probability 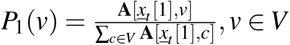. Then, we compute the “probability of acceptance” *β*, and draw a value *u* ∈ [0,1] uniformly at random. If *u* > *β*, then we accept *<u>x</u>*_(*t*+1)_ [1] = *v*, otherwise we reject *v* and set 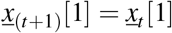. From *<u>x</u>*_(*t*+1)_[1] we perform a *k* – 1 step random walk to generate *<u>x</u>*(*t*+1)[2] to *<u>x</u>*_(*t*+1)_ [*k*] (for details, see the Supplement Section 5.2 Algorithm 2).

**Figure 3.**
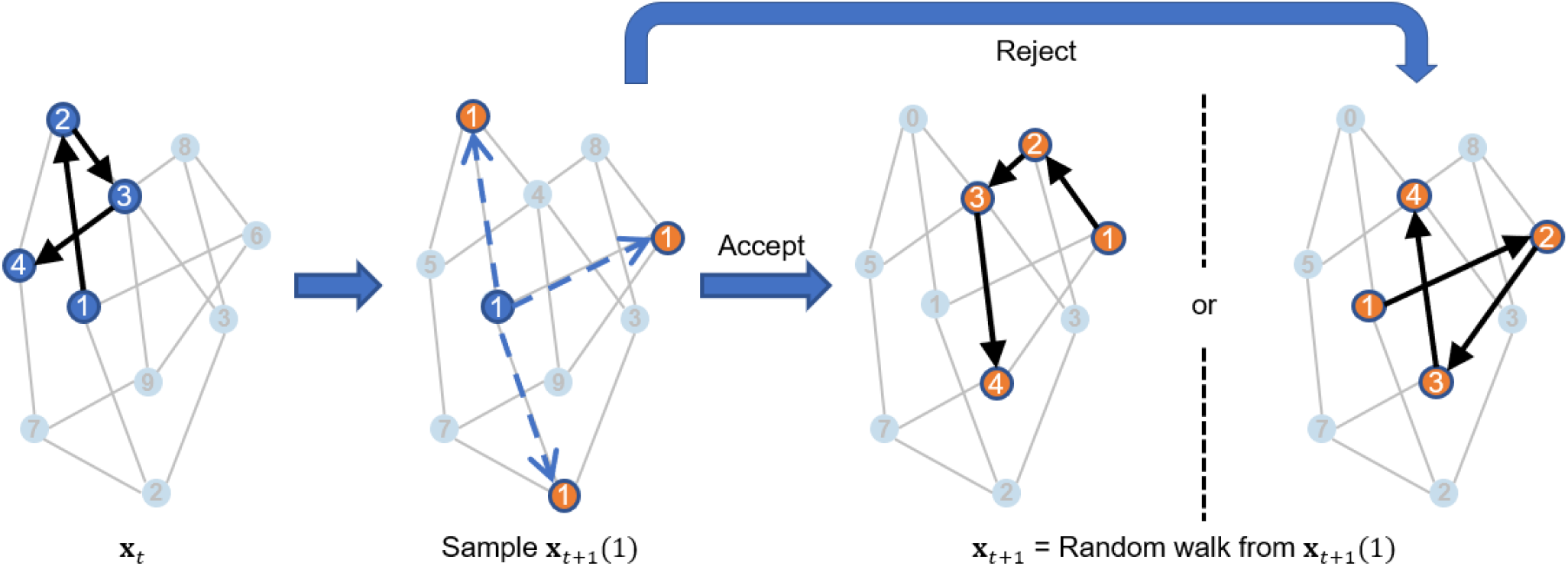
Workflow of the MCMC sampling algorithm, explaining how to generate from the sample at time t, *<u>x</u>_t_*, a sample at time (*t* + 1), *<u>x</u>*_(*t*+1)_.

**Algorithm 2.**
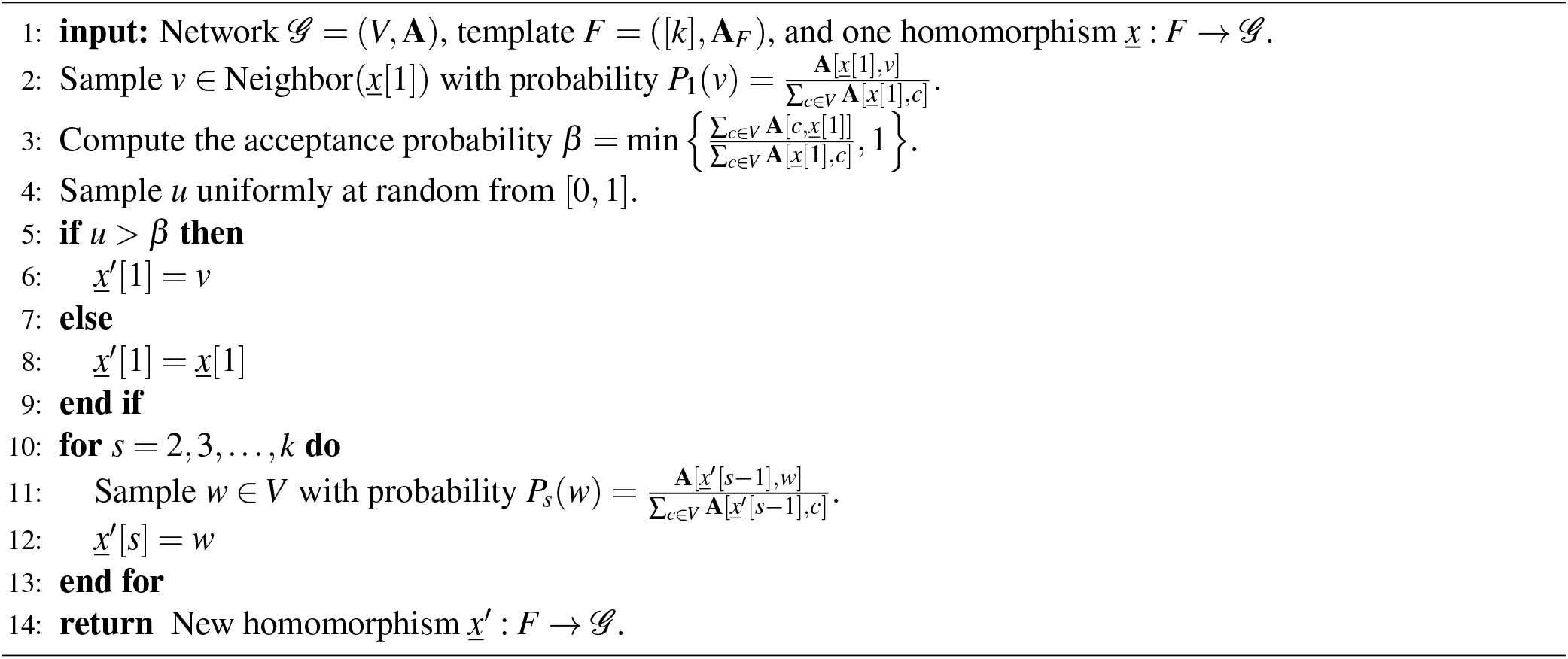
The MCMC Sampling Algorithm.

### Online convex NDL (online cvxNDL)

We start by initializing the dictionary **D**_0_ and representative sets 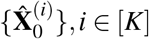, for each dictionary element (see the Supplement Section 5.2 Algorithm 3). After initialization, we perform iterative optimization to generate **D**_*t*_ and 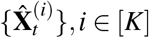, to reduce the loss at round *t*. At each iteration, we use MCMC sampling to obtain a *k*-node random subnetwork as sample **X**_*t*_, and then update the codes 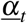 based on the dictionary **D**_*t*-1_ by solving the optimization problem in Equation (2). Then we assign the current sample to a representative set of the closest dictionary element, say **D**_*t*-1_[:, *j*], and then jointly update its representative set 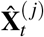 and all dictionaries **D**_*t*_ as shown in Figure 1 (see the Supplement Section 5.2 Algorithm 4).

**Algorithm 3.**
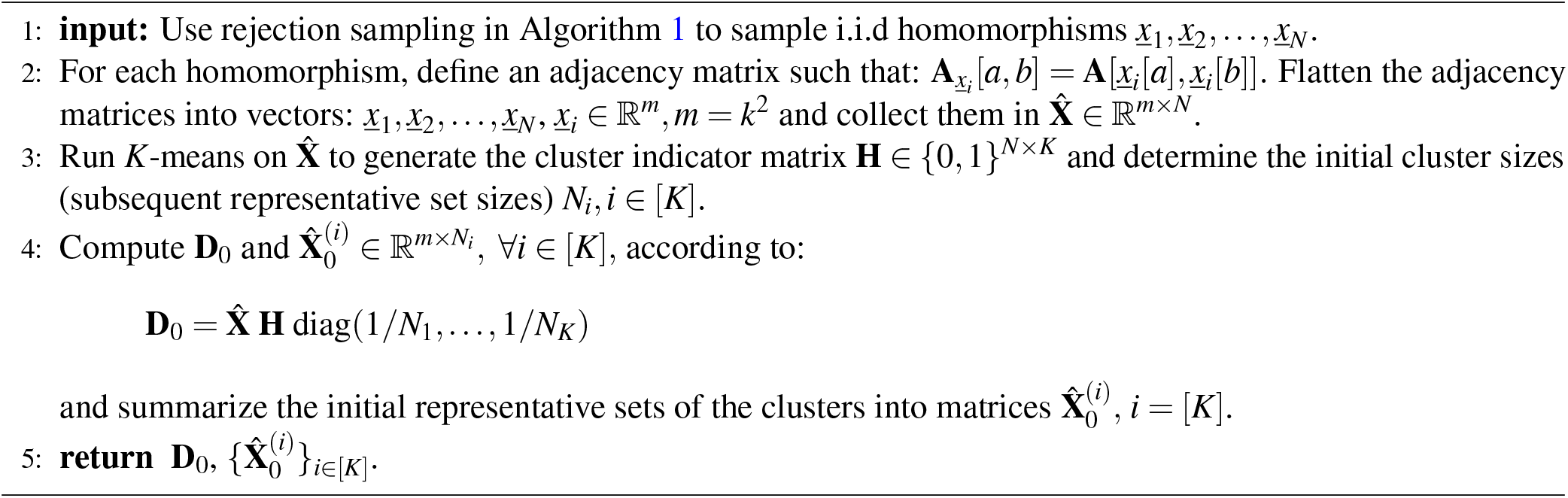
Initialization.

**Algorithm 4.**
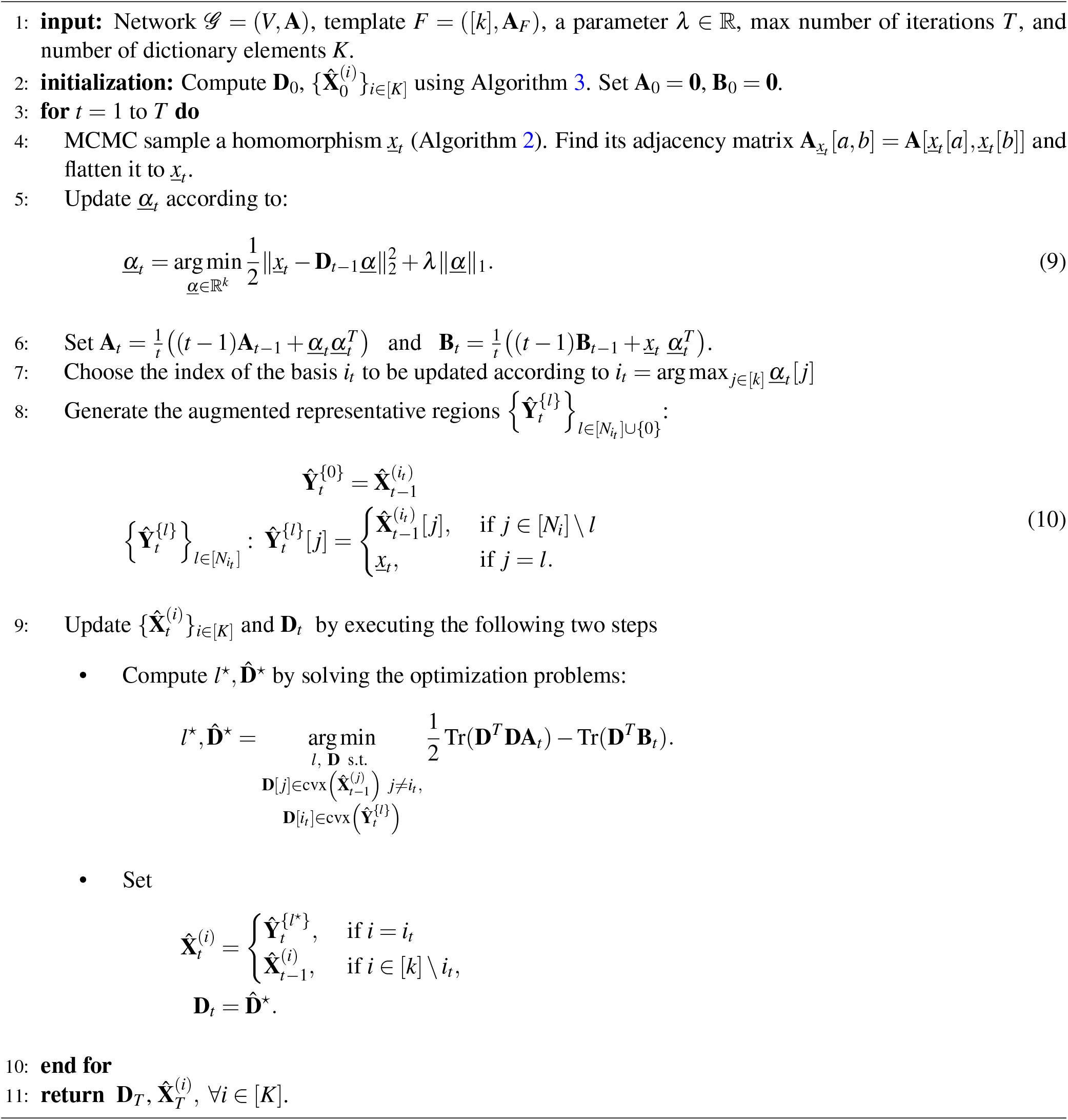
Online cvxNDL.

The output of the algorithm is a dictionary matrix **D**_*T*_ ∈ ℝ^*k*2×*K*^, where each column is a flattened vector of a dictionary element of size *k × k*, and the representative sets 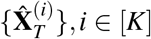, for each dictionary element. Each representative set 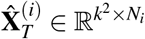 contains *N_i_* history-sampled subnetworks from the original network as its columns. We can can easily convert both the dictionary elements and representatives back to *k × k* adjacency matrices. Due to the added convexity constraint, each dictionary element **D**_*T*_ [:, *j*] at the final step *T* has the *interpretable form*:

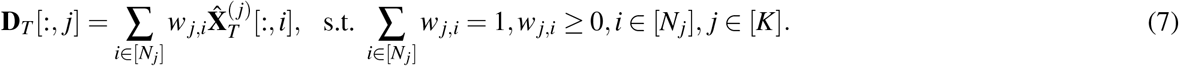

The weight *w_j,i_, i* ∈ [*N_j_*] is the *convex coefficient* of the *i*^th^ representative of dictionary element **D_T_**[:, *j*]. Dictionary elements learned from the data stream can be used to reconstruct the original network by multiplying it with the code *<u>α</u>_T_* obtained from Equation (2). The *j*^th^ index of *<u>α</u>*_*T*_ correspond to the contribution of dictionary element **D**_*T*_ — [:, *j*] to the reconstruction. Similarly to^29^, we can also define the *importance score* for each dictionary element as:

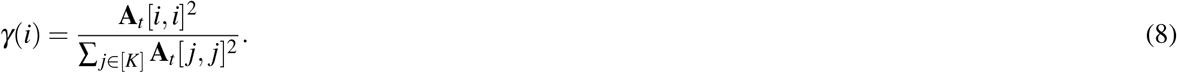

## 3 Results

We applied our online cvxNDL algorithm on both synthetic dataset and the ChIA-Drop data from *Drosophila Melanogaster* S2 cells on a dm3 reference genome (see Figure 5 for an illustration of the ChIA-Drop pipeline). Due to space limitations, the results pertaining to synthetic data and parts of the real data are deferred to the Supplement Section 5.3-5.5.

**Figure 4.**
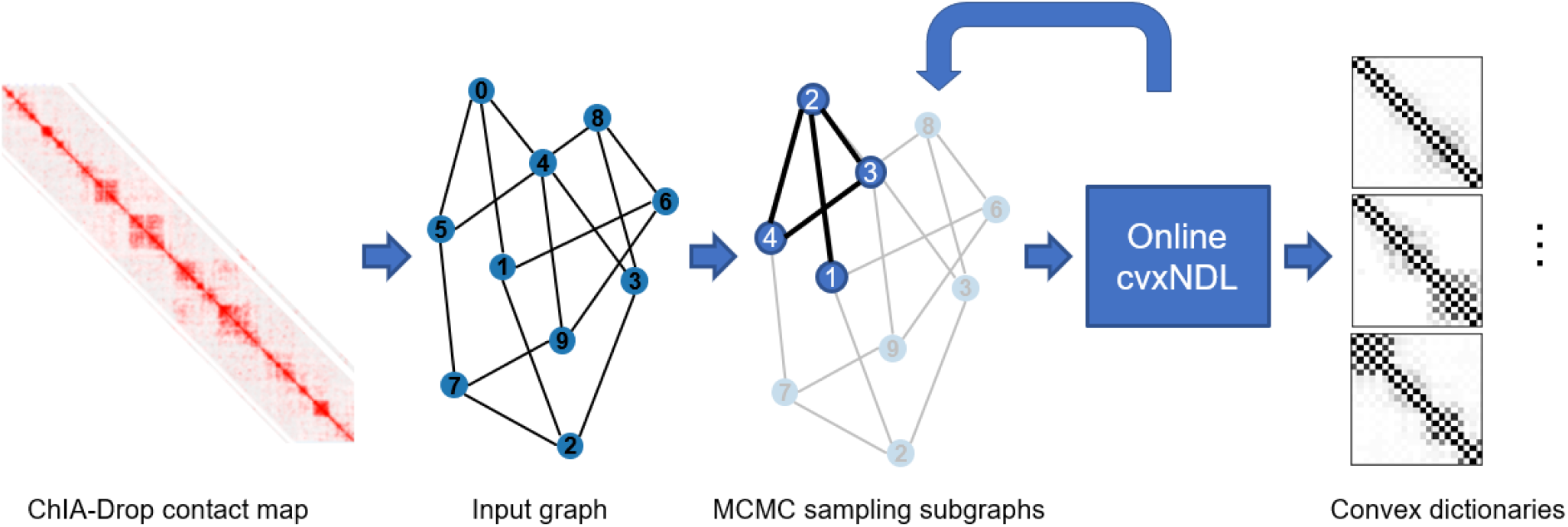
Workflow of online cvxNDL.

**Figure 5.**
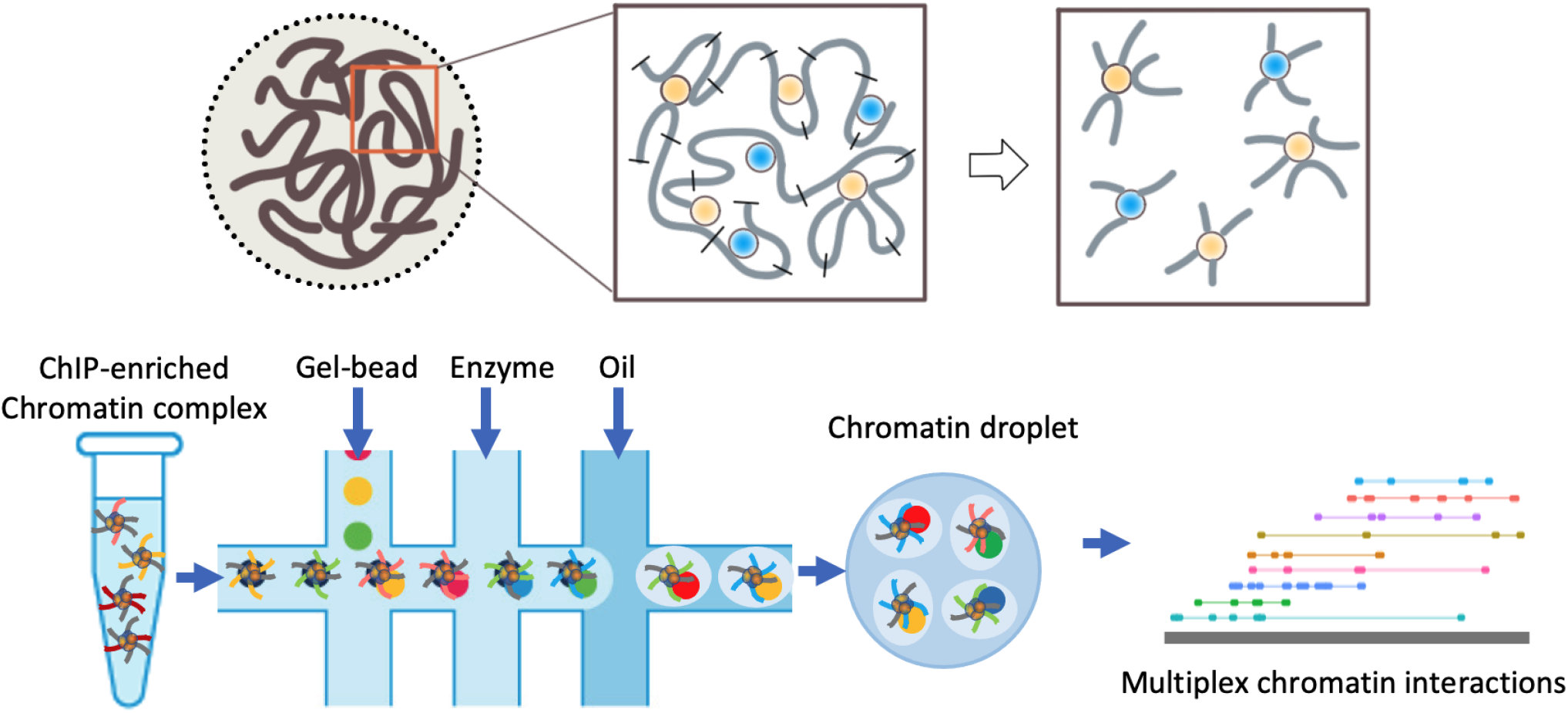
Generation of ChIA-Drop data. Chromatin samples are linked and fragmented (1^st^ row), guided to a microfluidic device for sequencing (2^nd^ row); reads are mapped to the reference to identify interaction complexes.

For analysis, we grouped 500 consecutive bases to form vertices for each chromosome. The ChIA-Drop data comprises information from chromosomes chr2L, chr2R, chr3L, chr3R, chr4 and chrX. Since chr4 and chrX are relatively small and most of the functional genes are located in chr2L, chr2R, chr3L, and chr3R, we focused our experiments on the remaining four chromosomes. To create the network, we converted the multiway interactions (i.e., hyperedge measurements) into cliques, following the well-established protocol of *clique expansion* for generating networks from hypernetworks^32,33^. For all experiments, we set the number of dictionary elements to *K* = 25, and used template subnetworks of the form of paths, each including 21 nodes (e.g., 21 × 500 bases). The initialization step involves MCMC sampling of 500 subnetworks from the networks obtained as above, so that each dictionary element has at least 10 representatives. The maximum number of online steps (i.e., samples) is set to 1 million (see also Figure 4). Note that our algorithm is the first method for online learning of convex (interpretable) network dictionaries, and the ground truth dictionaries are not known as they are also being studied for the first time. We therefore compare online cvxNDL with NMF, CMF and online NDL, but only in terms of global network reconstruction accuracy using the derived dictionaries. More comprehensive results are reported in the Supplement Section 5.4.

As a consequence of the convexity constraint, every dictionary element has a set of representatives that corresponds to a real observed subnetwork whose nodes can be mapped back to actual genomic loci. This allows us to find genes that overlap with at least one node included in some representative. Using these “covering genes”, we run the GO enrichment analysis from http://geneontology.org under annotation category “Biological Process” and with the reference list *“Drosophila Melanogaster”* for each dictionary element. We selected results with false discovery rate (FDR) < 0.05 as our candidate set for enriched GO terms. Note that there may be some inherently enriched GO terms for each dictionary element due to the sampling bias, learned from samples coming from the same chromosome. To remove this bias, we ran another GO enrichment analysis with all genes in each chromosome and used that results to filter out the undesired background GO terms in each dictionary element.

We also used the hierarchical structure of GO terms^34^. There, GO terms represent nodes in a directed acyclic graph while arcs indicates their relationship. A child GO term is considered more specific than and parent GO term. But since a child node may have multiple parents nodes, we further process our results as follows. For each GO term, i) we first find all the paths between it and the root node (which is “Biological Process” in our setting), and ii) we remove all intermediate parent GO terms from its enriched GO terms set. By iteratively repeating this filtering process for each dictionary element, we arrive at a list of most specific GO terms for each dictionary element. For more details behind the GO enrichment analysis, see the Supplement Section 5.5.

### Discussion

The dictionary elements for Drosophila ChIA-Drop data, chr2L, chr2R, chr3R and chr3L, obtained via online cvxNDL are shown in Figure 6. A comparison of the dictionaries constructed using online cvxNDL and other methods for chr2L is shown in Figure 7. We color-coded each representative based on the actual genomic location of their cover genes; we also used the weights in the convex combination 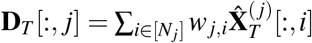 to color-code each dictionary element as well. Therefore, each dictionary element doe not only provide an interaction pattern but also captures the genomic locations involved, along with their importance factor. We also report the span of representatives, defined as the largest distance between genomic fragments (nodes) covered by the representative in Figure 8. The span captures the distances amongst interacting elements and the reported results are organized by chromosome. Corresponding results for the densities are reported in the Supplement Section 5.5.2. In comparison, classical NMF only captures partial structural patterns, and does not allow mapping back the results to actual genomic regions. It is hence hard to give a biological interpretation of each dictionary element; online cvxNDL and NDL both use a *k*-chain as the template. But while the dictionary elements obtained via CMF have large spreads, those generated by online cvxNDL have smaller yet still significant spreads that are likely to capture meaningful long-range interactions. Compared to online NDL, online cvxNDL also has a more balanced distribution of importance scores. For example, in Figure 7(b), dict_0 has score 0.459, while the scores in Figure 7(d) are all ≤ 0.085. Results for other chromosomes are in the Supplement Section 5.4.1.

**Figure 6.**
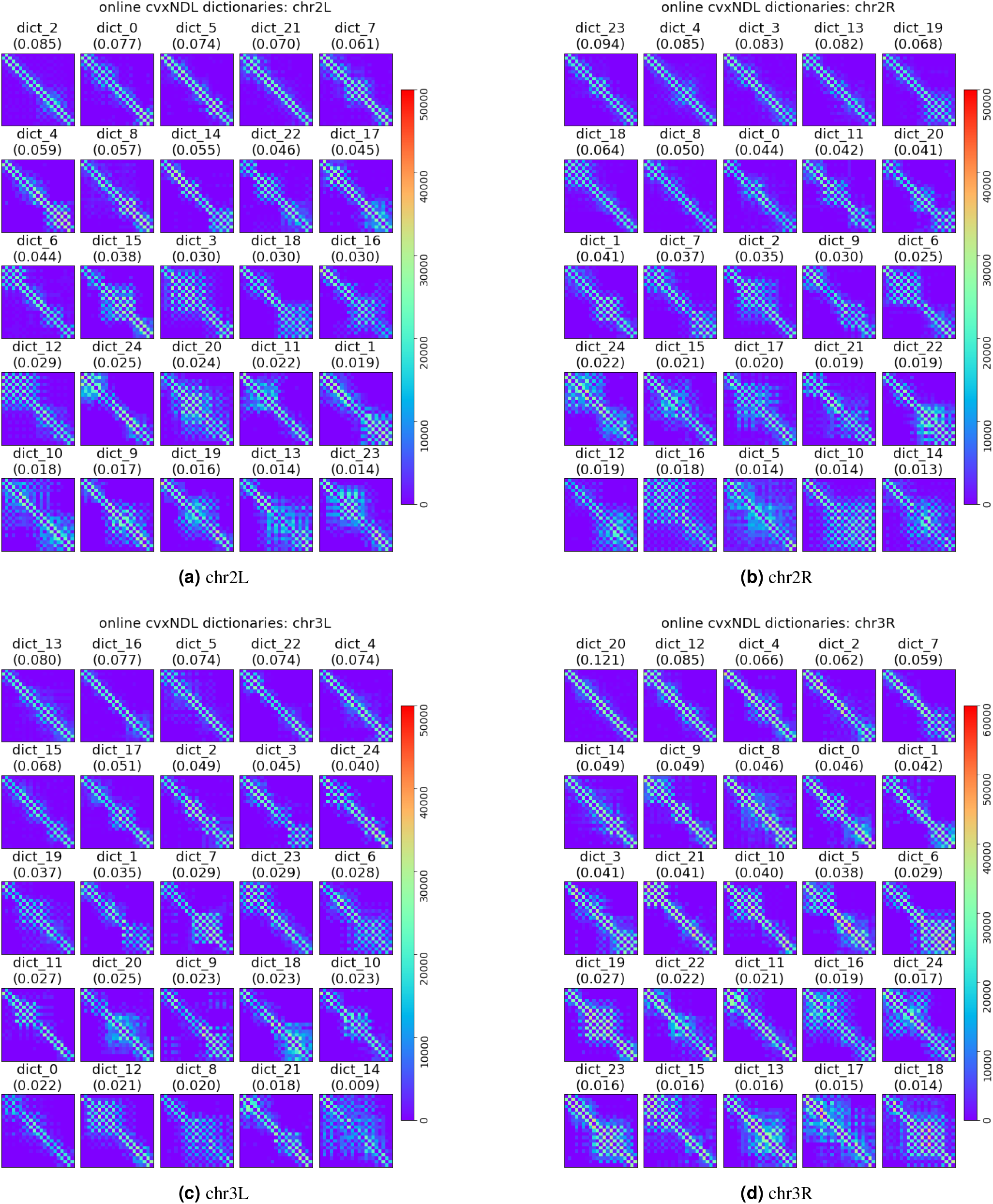
Dictionary elements for *Drosophila* chromosomes 2L, 2R, 3L and 3R obtained using online cvxNDL. Each subplot contains 25 dictionary elements for the corresponding chromosome and each block in the subplots corresponds to one dictionary element. The elements are ordered by their importance score.

**Figure 7.**
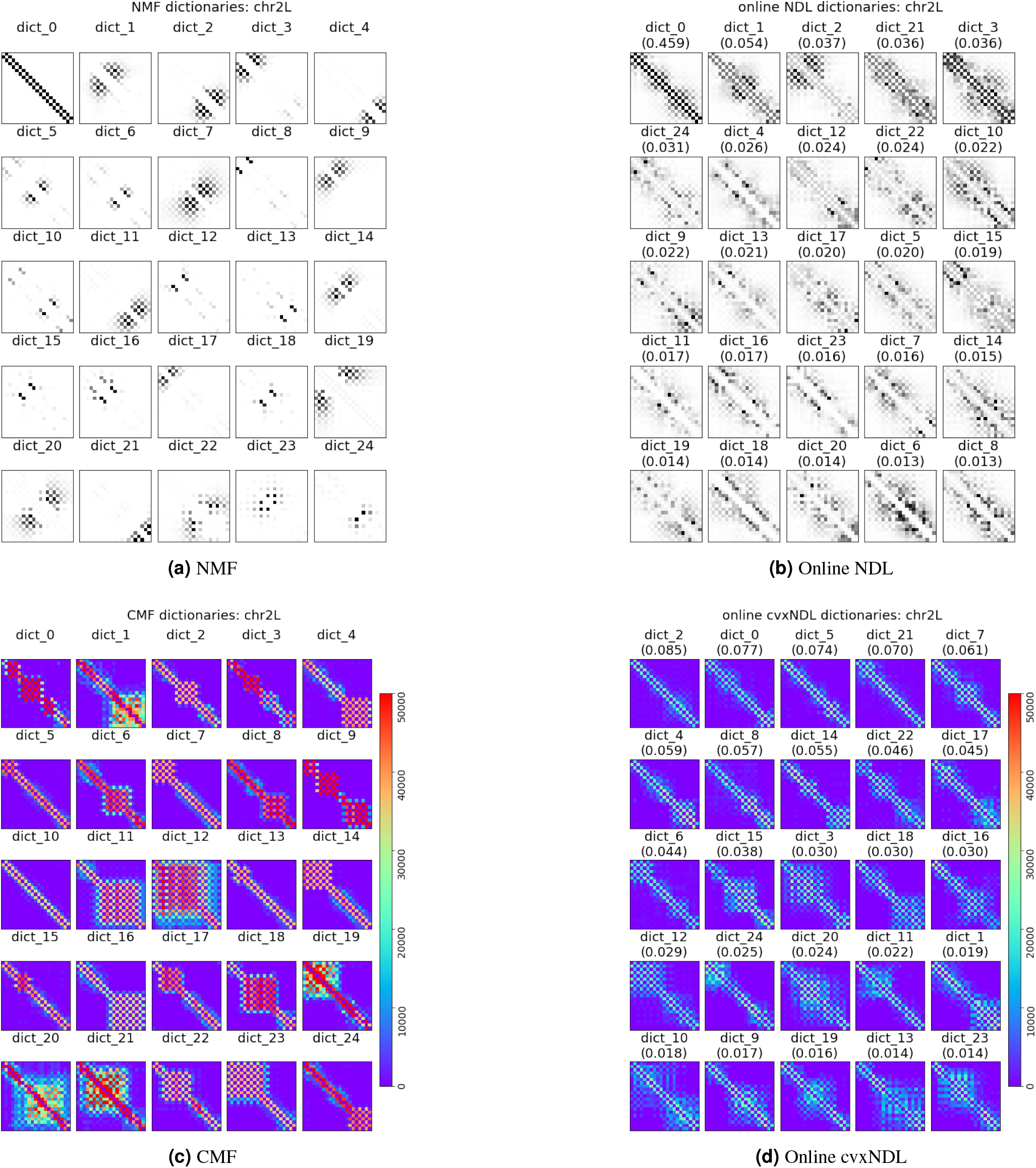
Dictionary elements for Drosophila chromosome chr2L generated by NMF (7a), online NDL (7b), CMF (7c) and online cvxNDL (7d). NMF and CMF are learned off-line, using a total of 20,000 samples. Note that these algorithms do not scale and cannot work with larger number of samples such as those used in online cvxNDL. The color-coding is performed in the same manner as for the accompanying online cvxNDL results. Columns of the dictionary elements in the second row are color-coded based on the genome locations of the representatives. As the locations can be determined only via convex methods, the top row for NMF and online NDL is black and white.

**Figure 8.**
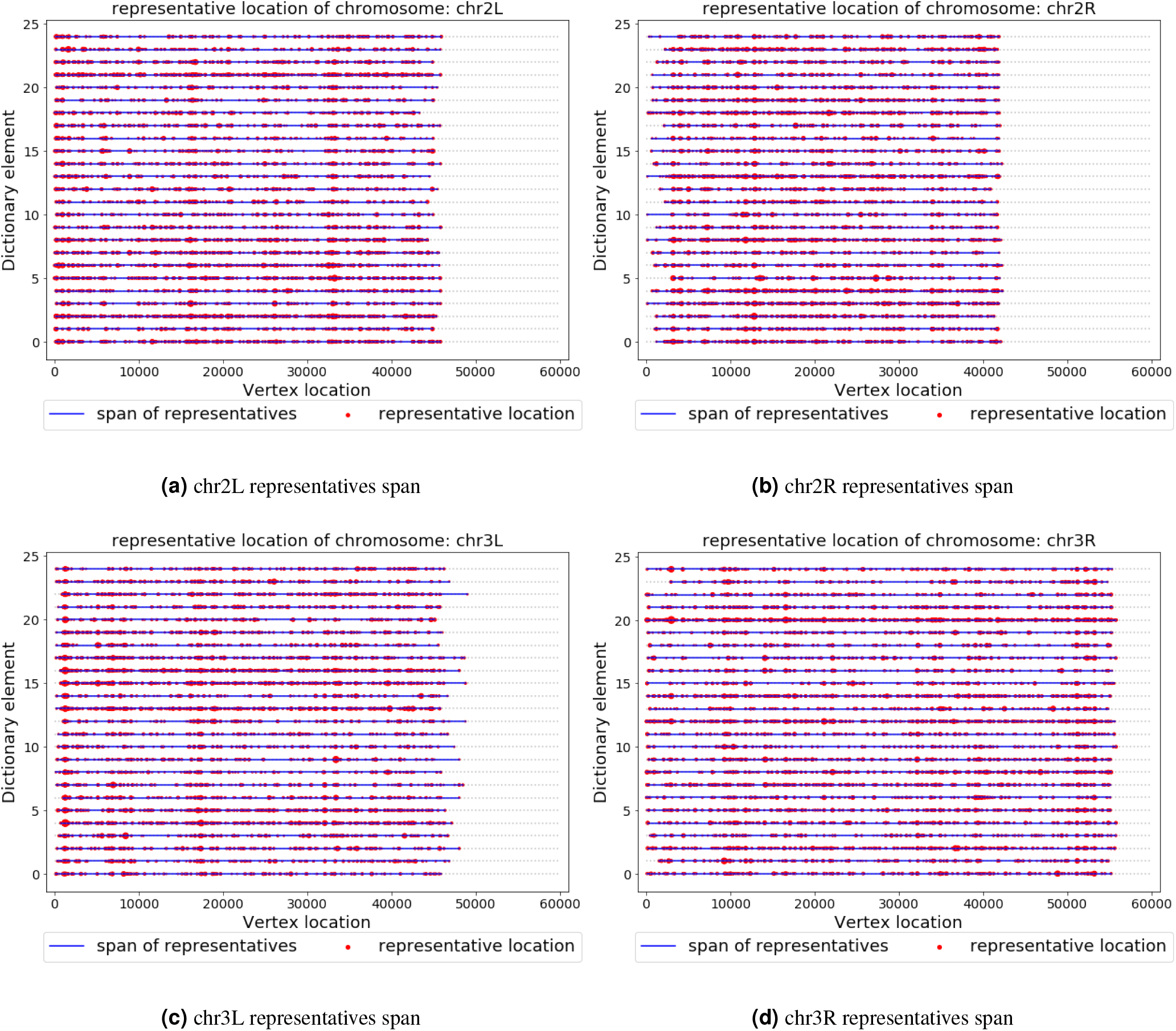
Span of the representatives. Blue lines correspond to chromatin lengths over which interactions are observed, while the red dots indicating the genome locations of nodes that appear in the representatives. The sizes of the red dots are indicative of the weight of the representative in the convex combination.

### Reconstruction accuracy

Once a dictionary is constructed, one can use the network reconstruction algorithm from^29^ to reconstruct a subnetwork or the whole network by locally approximating subnetworks via dictionary elements. Accuracy of approximation in this case measures the “expressibility” of the dictionary with respect to the network. All methods, excluding randomly generated dictionaries used for illustration only, can accurately reconstruct the input network. Only subtle and nearly insignificant visual differences are observable in off-diagonal connections representing TADs, which are shown in the Supplement Section 5.4.2. For a more quantitative assessment, the average precision recall score for all methods are plotted in Table 1, and a zoomed in sample based reconstruction result is shown in the Supplement Figure 16. As expected, random dictionaries have the lowest scores across all chromosomes, while all other methods are comparable. Hence, without reducing accuracy, online cvxNDL scales to large datasets due to its online nature and in addition allows for precise interpretation of the results. More details are provided in the Supplement Section 5.4.

**Table 1.** Average Precision Recall for different DL methods on all chromosomes and synthetic datasets, described in the Supplement Section 5.3-5.4.

|  | chr2L | chr2R | chr3L | chr3R | Synthetic |
| --- | --- | --- | --- | --- | --- |
| Online cvxNDL | 0.9954 | 0.9986 | 0.9830 | 0.9876 | 0.9747 |
| Online NDL | 0.9955 | 0.9986 | 0.9834 | 0.9880 | 0.9728 |
| NMF | 0.9952 | 0.9985 | 0.9829 | 0.9873 | 0.9774 |
| CMF | 0.9951 | 0.9985 | 0.9824 | 0.9870 | 0.9731 |
| Random Dict. | 0.0007 | 0.2547 | 0.5276 | 0.0796 | 0.1922 |

### GO enrichment results

For each chromosome, we counted how many GO terms are enriched in each of its dictionary elements. The results are reported in Table 2, along with the corresponding importance score of each dictionary element in parenthesis. We labeled the top 5 dictionary element with most enriched GO terms using bold font. The total number of enriched GO terms for each chromosome found by combining all dictionary elements, as well as the number of unique GO terms, are reported in the last two rows. From Table 2, one can see that dictionary elements with 0 enriched GO terms all have small importance scores (mostly below 0.04, and the largest possible value 0.059). Dictionary elements with higher importance scores all tend to involve a larger number of enriched GO terms. A detailed collection of tables describing the structure of each dictionary elements and its number of enriched GO terms can be found in the Supplement Section 5.5.2. We also report on the most frequently enriched GO terms and least frequently enriched GO terms on each chromosome, and present the corresponding dictionary elements which were found to be enriched. The most frequent GO terms were associated with regulatory functions, reflecting the role of RNA Polymerase II. Table 3 illustrates the findings for chr3R, while the results for other chromosomes are presented in the Supplement Section 5.5.1.

**Table 2.** Number of enriched GO terms for each dictionary element, along with the corresponding importance score (in brackets). The top-5 dictionary elements according to their number of enriched GO terms are labeled in boldface font for each chromosome. The last two rows show the total number of enriched GO terms and the number of unique enriched GO terms for each dictionary element, respectively.

|  | chr2R | chr2L | chr3L | chr3R |
| --- | --- | --- | --- | --- |
| dict_0 | 2 (0.077) | 4 (0.044) | 0 (0.022) | 15 (0.046) |
| dict_1 | 0 (0.019) | 1 (0.041) | 3 (0.035) | 9 (0.042) |
| dict_2 | **20 (0.085)** | 0 (0.035) | 0 (0.049) | 13 (0.062) |
| dict_3 | 0 (0.030) | **12 (0.083)** | 3 (0.045) | 7 (0.041) |
| dict_4 | 0 (0.059) | **40 (0.085)** | 3 (0.074) | **20 (0.066)** |
| dict_5 | 15 (0.074) | 0 (0.014) | **6 (0.074)** | 2 (0.038) |
| dict_6 | **19 (0.044)** | 0 (0.025) | 1 (0.028) | 2 (0.029) |
| dict_7 | **24 (0.061)** | 1 (0.037) | 1 (0.029) | 14 (0.059) |
| dict_8 | **31 (0.057)** | **17 (0.050)** | 0 (0.020) | **25 (0.046)** |
| dict_9 | 0 (0.017) | 0 (0.030) | 0 (0.023) | 1 (0.049) |
| dict_10 | 0 (0.018) | 0 (0.014) | 2 (0.023) | 5 (0.040) |
| dict_11 | 2 (0.022) | 1 (0.042) | 0 (0.027) | 0 (0.021) |
| dict_12 | 1 (0.029) | 2 (0.019) | 1 (0.021) | **16 (0.085)** |
| dict_13 | 0 (0.014) | 9 (0.082) | **16 (0.080)** | **57 (0.016)** |
| dict_14 | 6 (0.055) | 5 (0.013) | 0 (0.009) | 6 (0.049) |
| dict_15 | 0 (0.038) | **23 (0.021)** | **10 (0.068)** | 8 (0.016) |
| dict_16 | 2 (0.030) | 0 (0.018) | **14 (0.077)** | 0 (0.019) |
| dict_17 | 0 (0.045) | 0 (0.020) | **9 (0.051)** | 0 (0.015) |
| dict_18 | 0 (0.030) | 8 (0.064) | 4 (0.023) | 0 (0.014) |
| dict_19 | 0 (0.016) | 7 (0.068) | 0 (0.037) | 0 (0.027) |
| dict_20 | 0 (0.024) | 6 (0.041) | 0 (0.025) | **124 (0.121)** |
| dict_21 | **27 (0.070)** | 0 (0.019) | 0 (0.018) | 10 (0.041) |
| dict_22 | 1 (0.046) | 8 (0.019) | 4 (0.074) | 4 (0.022) |
| dict_23 | 0 (0.014) | **10 (0.094)** | 3 (0.029) | 0 (0.016) |
| dict_24 | 0 (0.025) | 2 (0.022) | 0 (0.040) | 4 (0.017) |
| Total # of GO terms | 150 | 156 | 80 | 342 |
| # of unique GO terms | 100 | 103 | 67 | 223 |

**Table 3.**
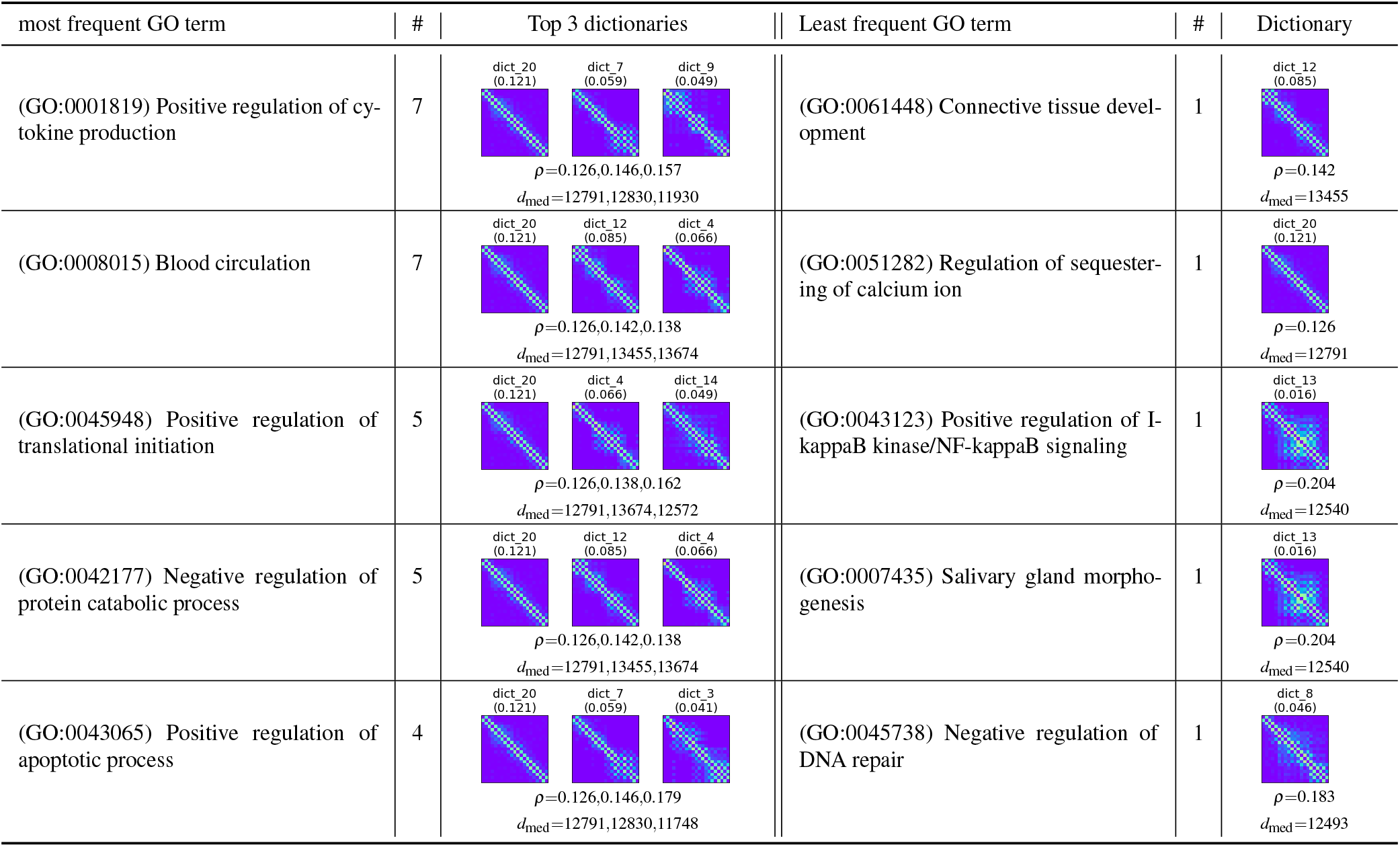
Top-5 enriched GO terms that occur most frequently and least frequently within the span of dictionary elements for chr3R. Column ‘#’ indicates the number of dictionary elements that show enrichment for the given GO term. Also we report up to 3 dictionary elements with largest importance score in the dictionary, along with the density of interactions in the dictionary element *ρ* and median distance of all adjacent pairs of nodes in its representatives *d*_med_.

| most frequent GO term | # | Top 3 dictionaries | Least frequent GO term | # | Dictionary |
| --- | --- | --- | --- | --- | --- |
| (GO:0001819) Positive regulation of cytokine production | 7 | dict_20<br>(0.121) dict_7<br>(0.059) dict_9<br>(0.049)<br>$\rho=0.126,0.146,0.157$<br>$d_{med}=12791,12830,11930$ | (GO:0061448) Connective tissue development | 1 | dict_12<br>(0.085)<br>$\rho=0.142$<br>$d_{med}=13455$ |
| (GO:0008015) Blood circulation | 7 | dict_20<br>(0.121) dict_12<br>(0.085) dict_4<br>(0.066)<br>$\rho=0.126,0.142,0.138$<br>$d_{med}=12791,13455,13674$ | (GO:0051282) Regulation of sequestering of calcium ion | 1 | dict_20<br>(0.121)<br>$\rho=0.126$<br>$d_{med}=12791$ |
| (GO:0045948) Positive regulation of translational initiation | 5 | dict_20<br>(0.121) dict_4<br>(0.066) dict_14<br>(0.049)<br>$\rho=0.126,0.138,0.162$<br>$d_{med}=12791,13674,12572$ | (GO:0043123) Positive regulation of I-kappaB kinase/NF-kappaB signaling | 1 | dict_13<br>(0.016)<br>$\rho=0.204$<br>$d_{med}=12540$ |
| (GO:0042177) Negative regulation of protein catabolic process | 5 | dict_20<br>(0.121) dict_12<br>(0.085) dict_4<br>(0.066)<br>$\rho=0.126,0.142,0.138$<br>$d_{med}=12791,13455,13674$ | (GO:0007435) Salivary gland morphogenesis | 1 | dict_13<br>(0.016)<br>$\rho=0.204$<br>$d_{med}=12540$ |
| (GO:0043065) Positive regulation of apoptotic process | 4 | dict_20<br>(0.121) dict_7<br>(0.059) dict_3<br>(0.041)<br>$\rho=0.126,0.146,0.179$<br>$d_{med}=12791,12830,11748$ | (GO:0045738) Negative regulation of DNA repair | 1 | dict_8<br>(0.046)<br>$\rho=0.183$<br>$d_{med}=12493$ |

## 4 Conclusion

We proposed a new online convex network dictionary learning algorithm for analyzing complex chromatin interaction patterns in ChIA-Drop data. Combining efficient MCMC sampling algorithms with alternative optimization constrained by convexity conditions, we implemented an online cvxNDL method that offers biological interpretability not possible by that of standard NDL. We also performed GO enrichment analysis that uses filtering based on a GO hierarchy. The proposed learning method can produce network dictionaries that i) accurately capture the topological patterns of short- and long-range interactions in the chromatin input network; ii) accurately reconstruct the original network using as few as 25 dictionary elements; and iii) have biologically interpretable meaning through their GO terms associated with each dictionary element.

## Funding and Acknowledgement

The work was supported by the National Science Foundation [1956384] and partially supported by National Science Foundation [2206296]. The authors gratefully acknowledge useful discussions with Dr. Yijun Ruan.

## 5 Supplement

### 5.1 Motivation

Although dictionary learning (DL), a form of (nonnegative) matrix factorization (MF), has been widely used in the analysis of biological data, *effective, efficient* and *biologically interpretable* computational methods for analyzing long-distance multiplexed chromatin interactions at a single-cell level are still lacking. This is mainly because most of the classical DL methods are not designed for network data. Furthermore, these interactions cannot be easily visualized or predicted via classical clustering approaches. This issue is best illustrated by Figure 9, where a part of the contact graph contains three hidden clusters, colored red, green, and blue^35^. When using a linear chromatin order, the particular structure of the clusters is not observable. By rearranging the rows/columns, the cluster structure becomes apparent within the adjacency matrix. To mitigate this issue, we propose a novel online convex network dictionary learning algorithm (online cvxNDL) and imposes “convexity” constraints on the sampled subgraph patterns to address both the issue of interpretability and scaling for graph-structured data. The approach and accompanying algorithmic implementations are described in the next section.

**Figure 9.**
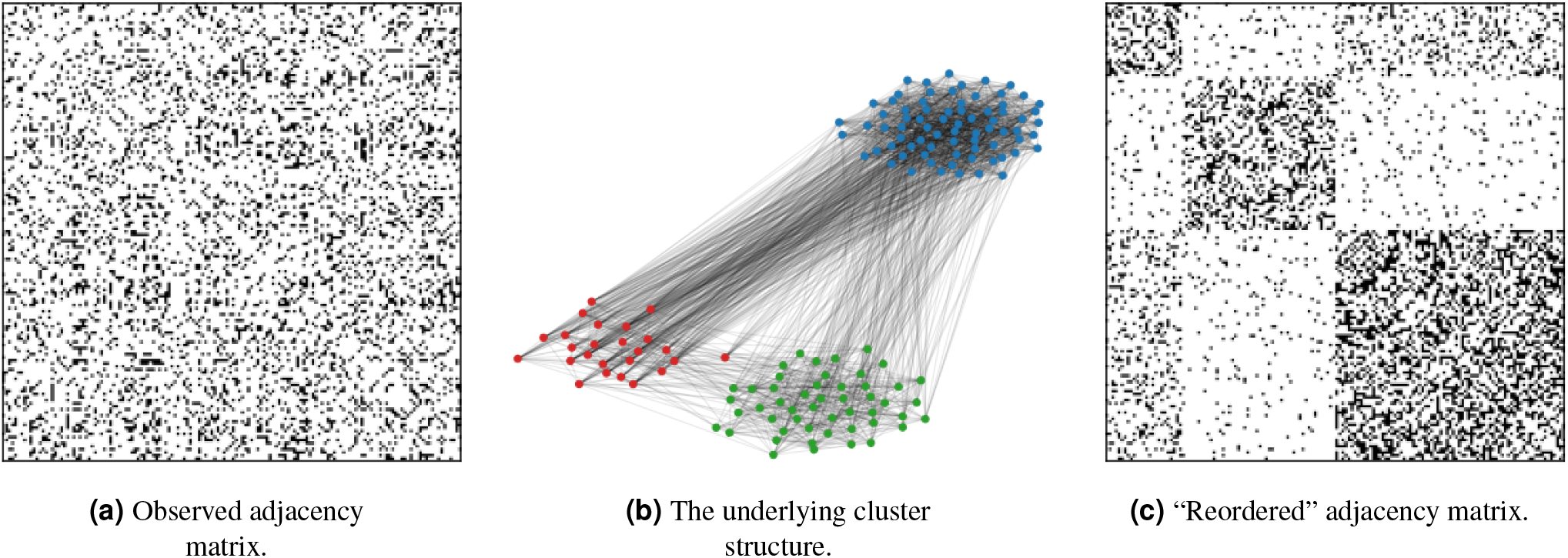
**(a)** Adjacency matrix of a three-cluster model, where points are arranged in linear order with dense interactions existing both at short and long-range. **(b)** The hidden cluster structure. **(c)** The reordered adjacency matrix that reveals all interaction classes.

### 5.2 Algorithmic Details

The algorithms presented in this sections describe the detailed steps of the implementations outlined in the Methods portion (Section 2) of the main text.

#### 5.2.1 MCMC Sampling of Subnetworks

The MCMC sampling algorithm has the goal to generate (sample) subnetworks induced by *k* nodes in the original input network 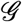, with the constraint that the subnetwork contains the template *F* topology. Note that one set of homomorphisms is defined as a vector of the form (with the assumption that 0^0^ = 1):

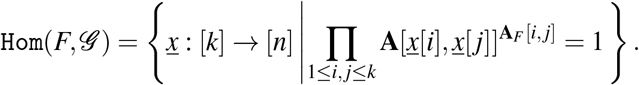

Algorithm 1 outlined how to use rejection sampling to obtain one homomorphism *<u>x</u>* (an illustrative example is presented in Figure 2). In this work, the choice of template network is a *k*-chain, a directed path from node 1 to k; chains are a simple and natural choice for networks that inherently contain long paths, such as chromatin interaction networks (since most measure contacts are due to proximity in the linear chromosome order).

Although one can find different homomorphisms from the input 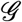 by repeatedly running Algorithm 1, this approach is computationally expensive. To efficiently generate a sequence of sample adjacency matrices 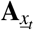 from *G*, the MCMC sampling algorithm gradually changes the sampled subnetwork based on previous samples as described in Algorithm 2. An illustrative example is shown in Figure 3. This sampling algorithm was introduced in^28,29^.

#### 5.2.2 Online Convex NDL (online cvxNDL)

Our online cvxNDL algorithm consists of two parts: Initialization and iterative optimization. For initialization (Algorithm 3), we need to compute an initial choice for the dictionary elements **D**_0_ and initialize the representative regions 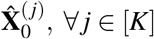. Note that we use i.i.d. sampling of homomorphisms only during the initialization step, and MCMC sampling afterwards. Upon initialization, we iteratively optimize the dictionary and the representative regions in the next phase (Algorithm 4). The output of the latter algorithm is the final dictionary **D**_*T*_ and the corresponding representative regions for all dictionary elements 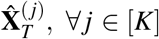. Due to the added convexity constraint, each dictionary element **D**_*T*_ [:, *j*] at the final step *T* has the following interpretable form:

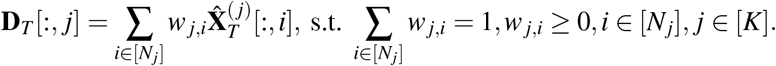

The weight *w_j,i_, i* ∈ [*N_j_*] is the convex coefficient of the *i*^th^ representative of dictionary element **D**_*T*_ [:, *j*].

### 5.3 Synthetic Data Analysis

We tested our online cvxNDL method on a network (graph) generated by Stochastic Block Model (SBM)^35^, containing 150 nodes with 3 clusters of size 25,50,75. Due to the small size of the synthetic set, we fixed the number of dictionary elements to *K* = 6, and used a chain of length 11 as our template. In the initialization step we sampled (collected) 30 subgraphs of the synthetic data, with each dictionary element represented by at least 3 representatives. The maximum number of iterations of the online method was set to 1, 000.

We compared online cvxNDL with various baseline methods, including NMF, CMF and online NDL. The learned dictionary elements for different methods are shown in Figure 10. The dictionary elements in online NDL and online cvxNDL are ordered by their importance score defined as 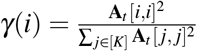. Each square block in the subplots indicates one dictionary element in the form of an adjacency matrix. The color-shade reflects the values in the adjacency matrix, with black corresponding to 1 (the largest value) and white corresponding to 0 (the smallest value).

**Figure 10.**
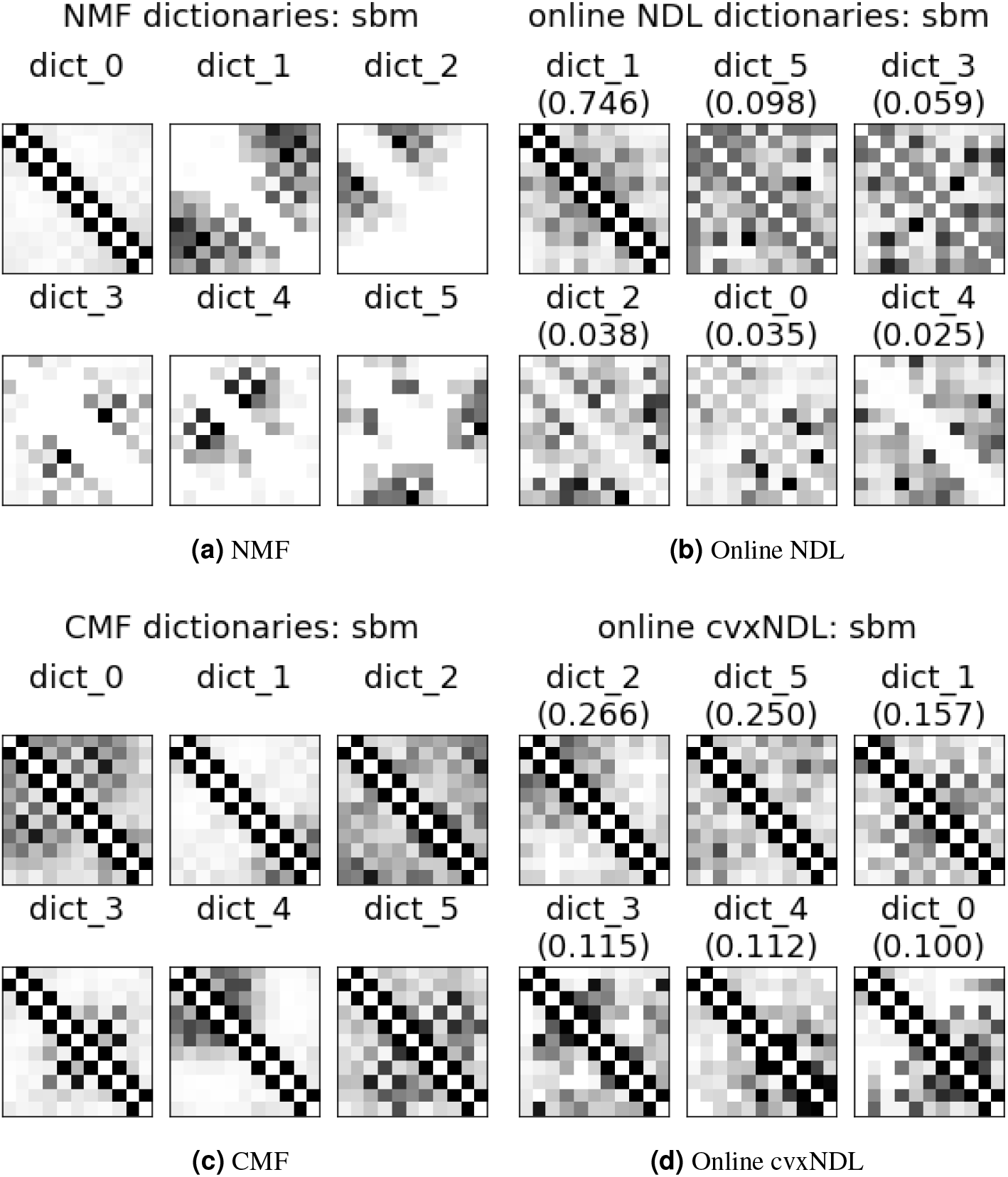
Dictionary elements generated by different MF methods on an SBM synthetic dataset. Numbers in parenthesis are the importance scores for online NDL and online cvxNDL.

From the results we can see that dictionaries generated using NMF only contains partial interaction structures and are hard to interpret. The two convex methods, CMF and online cvxNDL, contain the template structure in all learned dictionary elements, and show stronger off diagonal connectivity, which is expected as the input data has slightly stronger connections between the first and last cluster than other pairs (See Figure 11a). Online NDL dictionary elements represent “a middle ground” between NMF and online cvxNDL. Dictionary elements 2, 0 and 4 resemble those generated by NMF, while dictionary elements 1, 5 and 3 are similar to the ones generated by online cvxNDL, although with weaker connectivity. Also, the importance score distributions of online NDL and online cvxNDL differ substantial. In online NDL, dictionary element 1 in Figure 10b) represents the dominant component in representations, whereas in online cxvNDL, the top two dictionary elements (dictionary elements 2 and 5 in 10d) share similar scores and the dictionary elements in general have a more balanced distribution of importance scores. From the original adjacency we can see that there are indeed two different connectivity patterns in the network captured by online cvxNDL.

**Figure 11.**
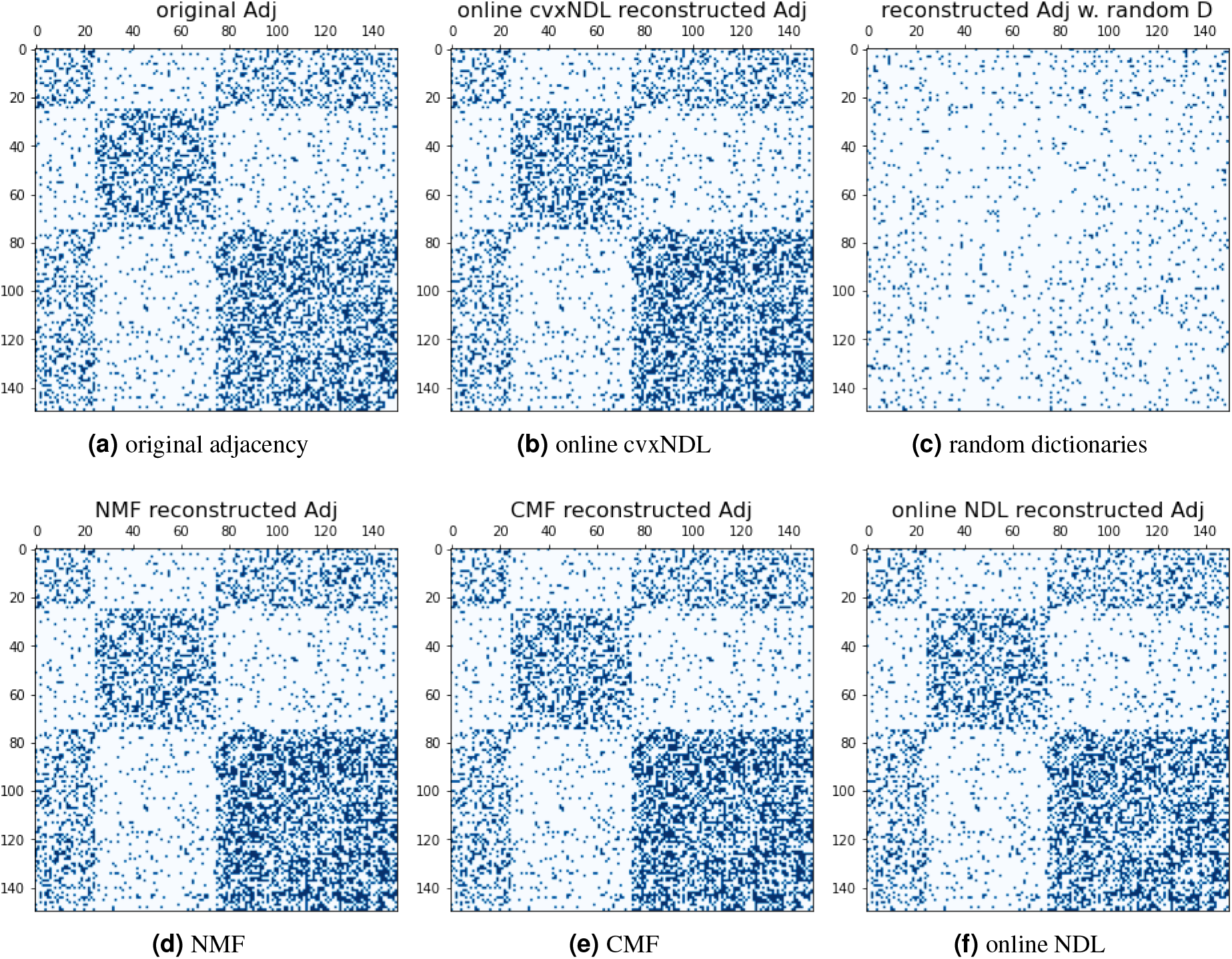
Original adjacency matrix and reconstructed adjacency matrices based on different DL methods and using random dictionaries.

**Reconstruction accuracy:** To validate the reliability of our learned dictionaries for representing the global interactions, we reconstructed the whole graph by aggregating the regenerated subgraphs: 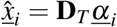 from the same MCMC sampling stream. For each method we selected the top-*m* edges after aggregation to reconstruct the original adjacency matrix, where *m* is the number of edges in the original adjacency matrix. The original and the reconstructed adjacency matrices are shown in Figure 11. For comparison, we also added the reconstructed adjacency achieved when using random dictionary elements. From the results we can see that all baseline methods, as well as online cvxNDL, almost perfectly reconstruct the original network, while, clearly random dictionaries do not capture any meaningful information. We also report the average precision recall score for each method, both for synthetic and real datasets as listed in Table 4.

**Table 4.** Average Precision Recall for different DL methods for all chromosome and the SBM synthetic dataset.

|  | chr2L | chr2R | chr3L | chr3R | Synthetic |
| --- | --- | --- | --- | --- | --- |
| Online cvxNDL | 0.9954 | 0.9986 | 0.9830 | 0.9876 | 0.9747 |
| Online NDL | 0.9955 | 0.9986 | 0.9834 | 0.9880 | 0.9728 |
| NMF | 0.9952 | 0.9985 | 0.9829 | 0.9873 | 0.9774 |
| CMF | 0.9951 | 0.9985 | 0.9824 | 0.9870 | 0.9731 |
| Rand. dictionaries | 0.0007 | 0.2547 | 0.5276 | 0.0796 | 0.1922 |

### 5.4 Results for Baseline Methods Applied to ChIA-Drop Datasets

#### 5.4.1 Dictionary Comparisons

Next, we describe dictionaries and reconstruction results for baseline methods on ChIA-Drop datasets corresponding to chromosomes chr2L, chr2R, chr3L, and chr3R. The results for online cvxNDL were reported in the main text.

#### 5.4.2 Reconstruction of ChIA-Drop Contact Maps

The reconstructions for 4 randomly selected subnetwork samples are shown in Figure 16, providing a means to visually assess the accuracy of reconstructed small-scale interactions.

**Figure 12.**
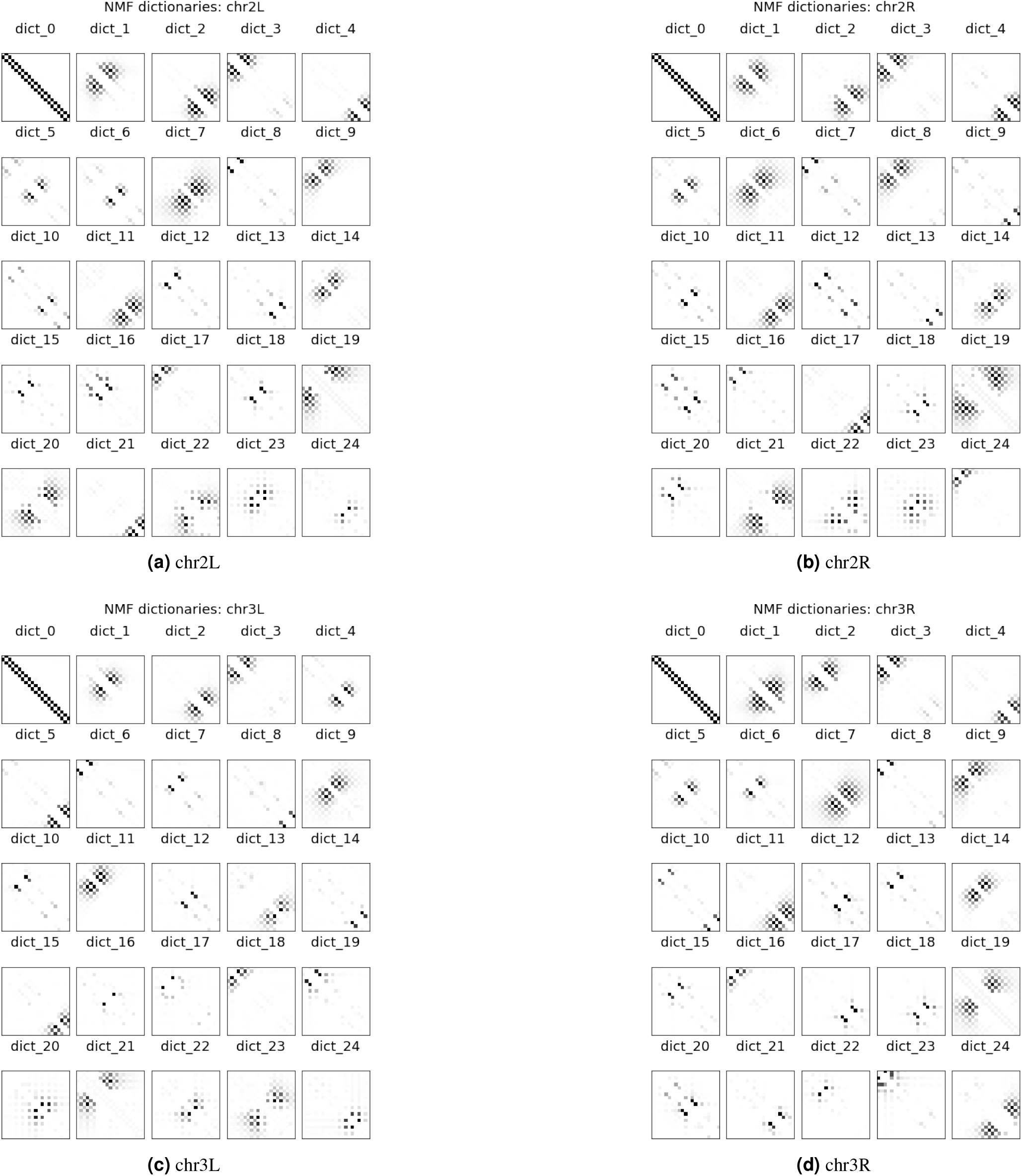
Dictionaries learned by NMF for chr2L, 2R, 3L and 3R.

**Figure 13.**
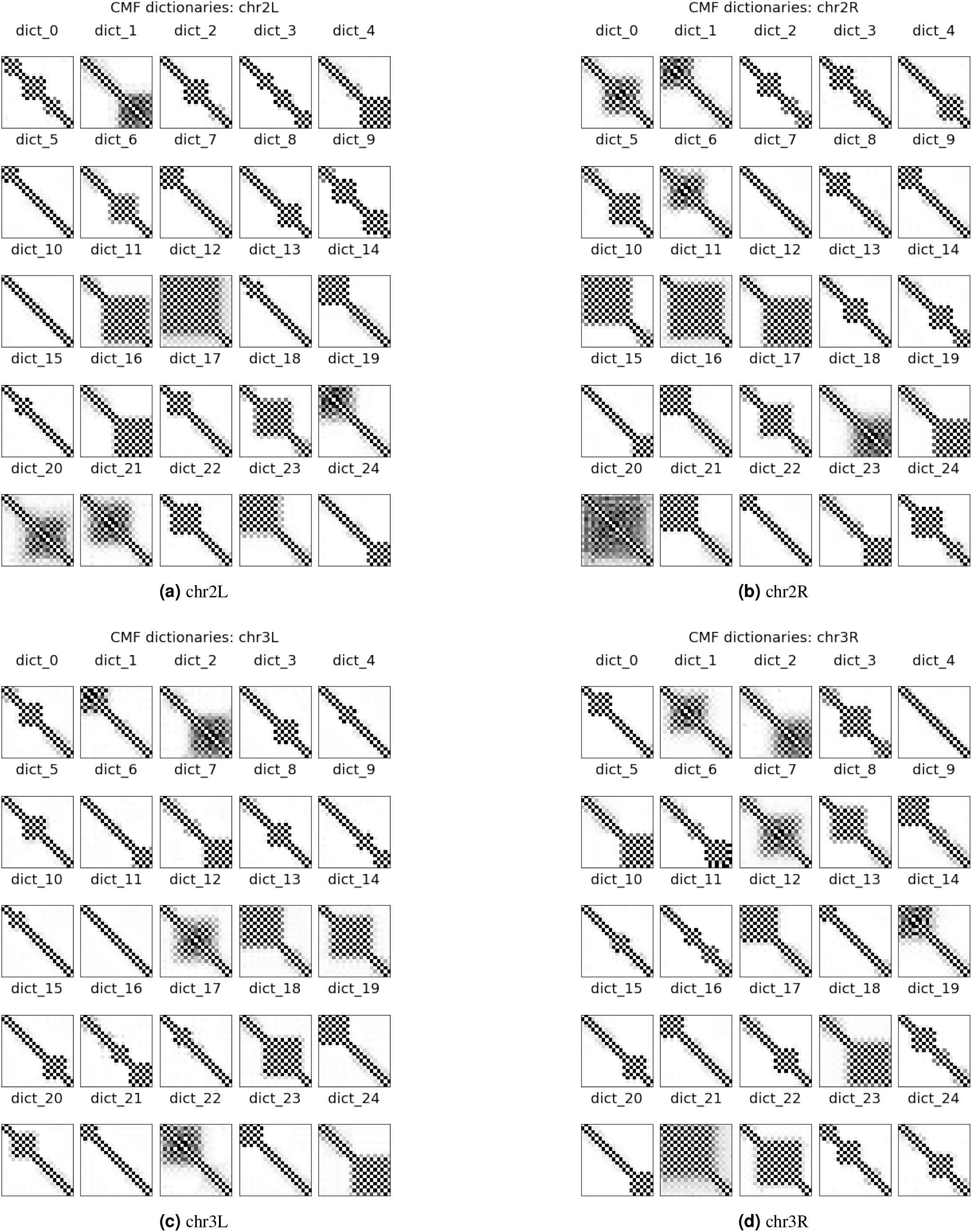
Dictionaries learned by CMF for chr2L, 2R, 3L and 3R.

**Figure 14.**
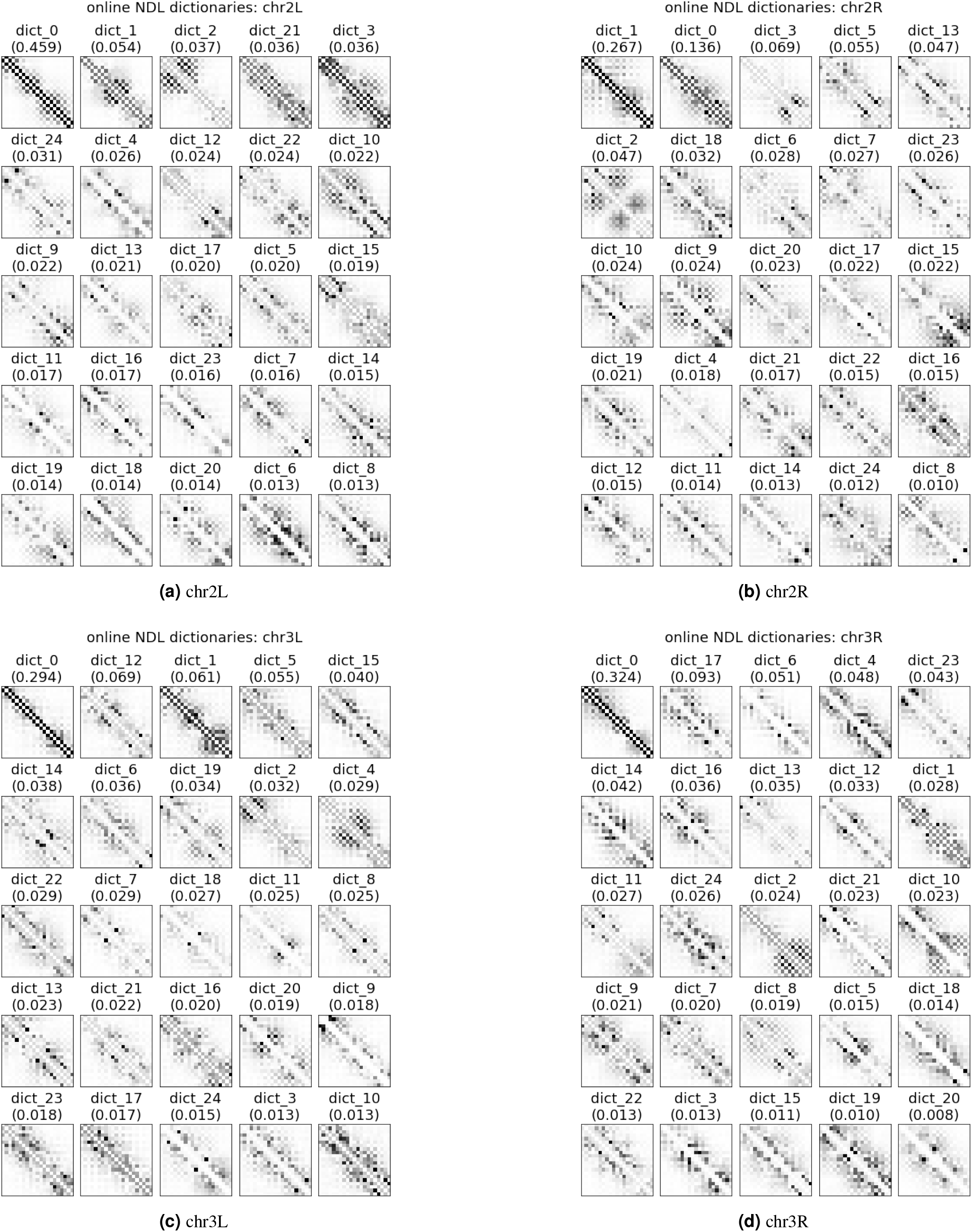
Dictionaries learned by online NDL for chr2L, 2R, 3L and 3R.

**Figure 15.**
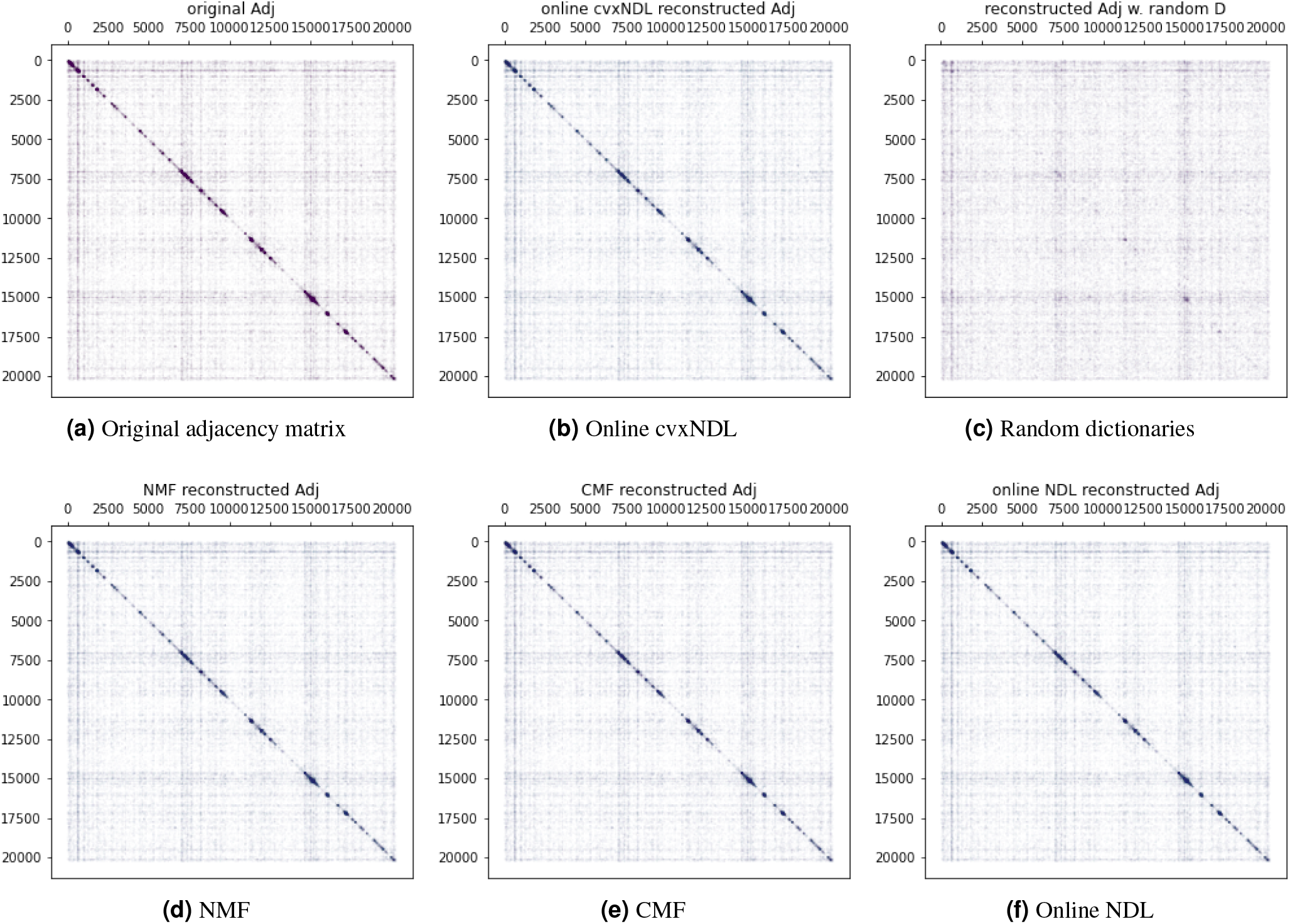
Comparison of network reconstructions obtained using different baseline methods and random dictionaries for Drosophila chromosome 2L (ChIA-Drop data). (a): The original adjacency matrix; (b, c, d, e, f): Reconstructed network adjacency matrices with online cxvNDL, random dictionary elements, NMF, CMF and online NDL, respectively.

**Figure 16.**
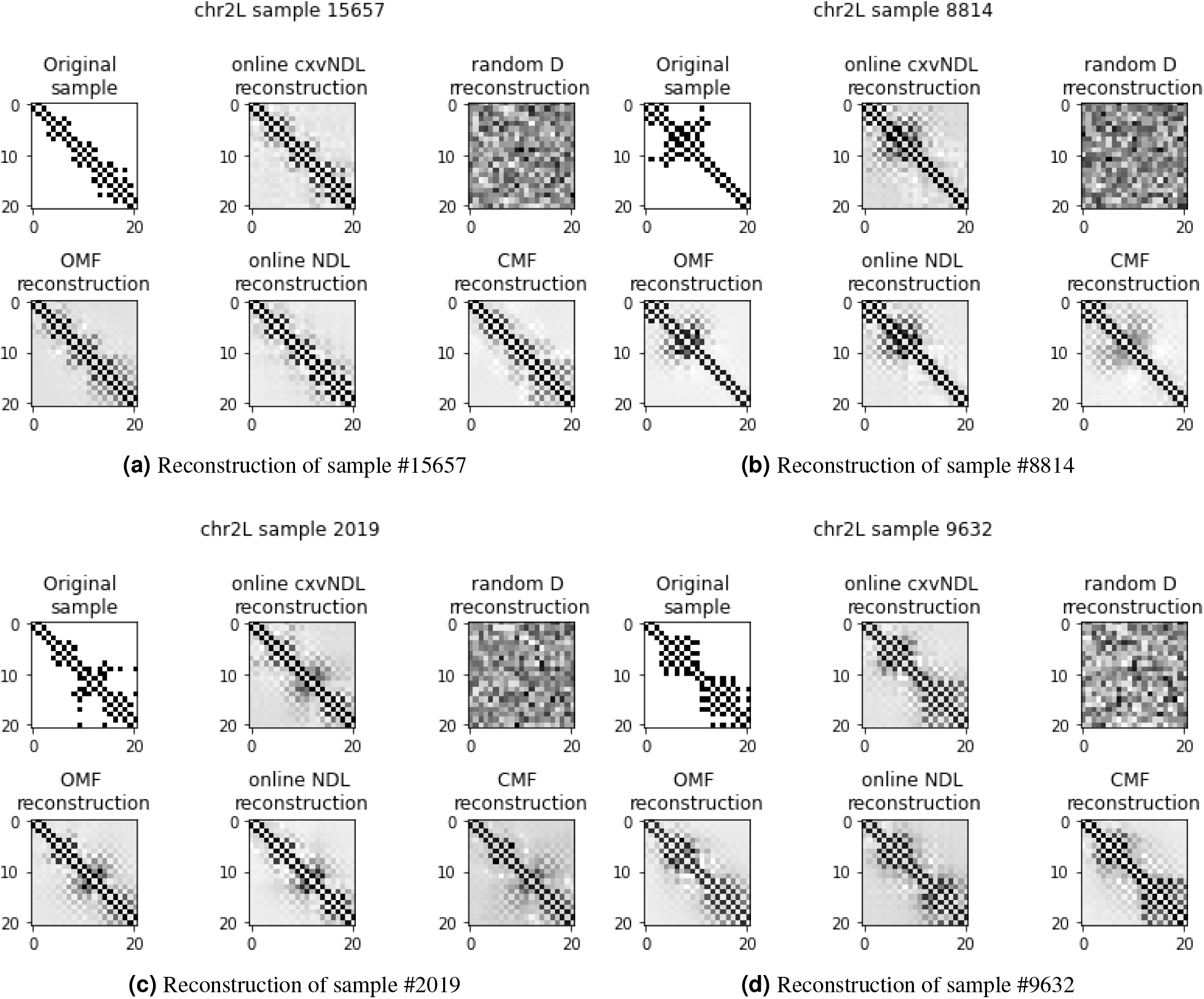
Reconstructed adjacency matrices for chr2L obtained using different methods and random dictionaries. OMF stands for Ordinary (Standard) MF.

**Figure 17.**
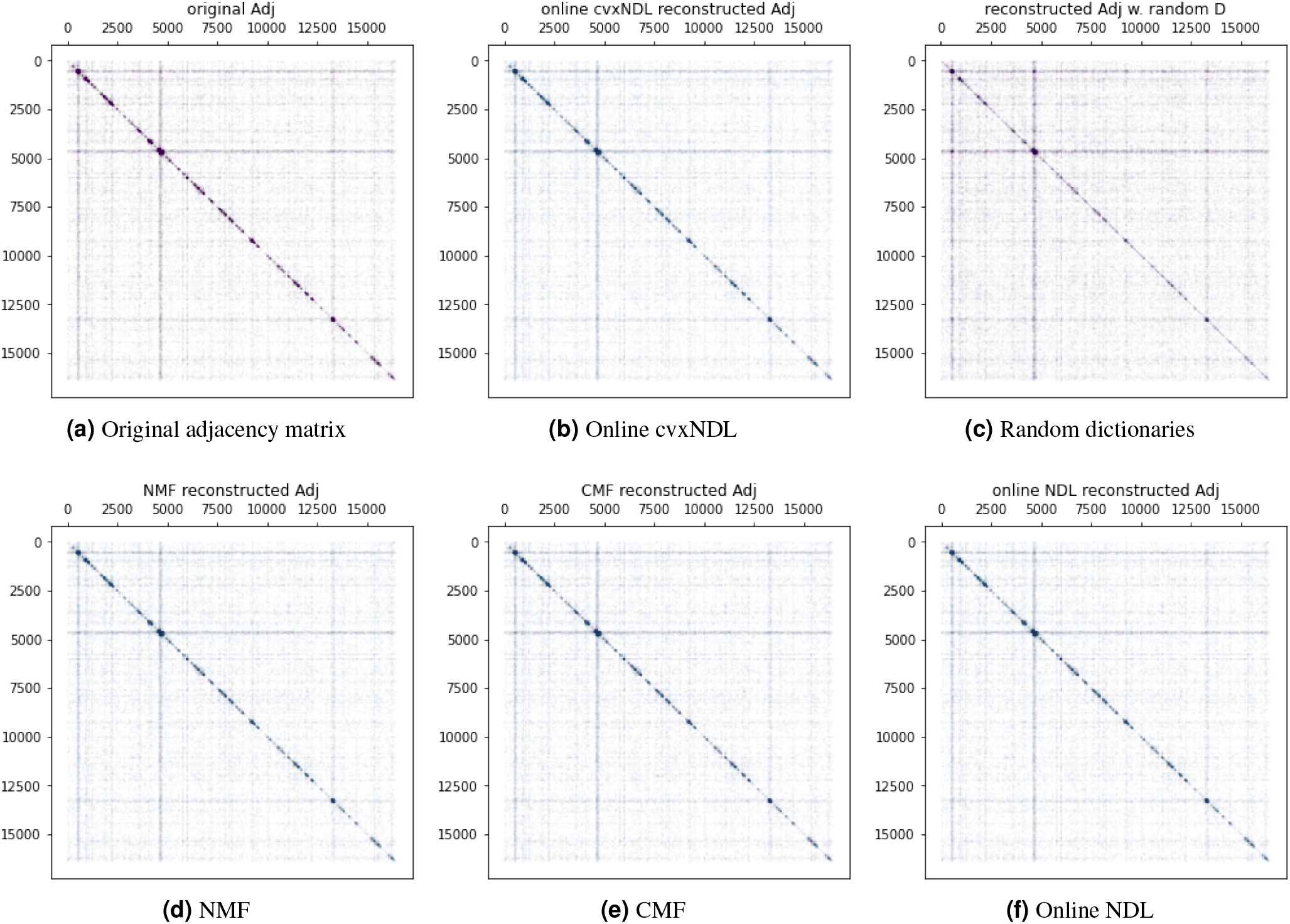
Reconstructed network comparisons based on different baseline methods and random dictionaries, applied on Drosophila chromosome 2R ChIA-Drop Data. (a): The original adjacency matrix. (b, c, d, e, f): Reconstructed network adjacency matrices with online cxvNDL, random dictionary elements, NMF, CMF and online NDL.

**Figure 18.**
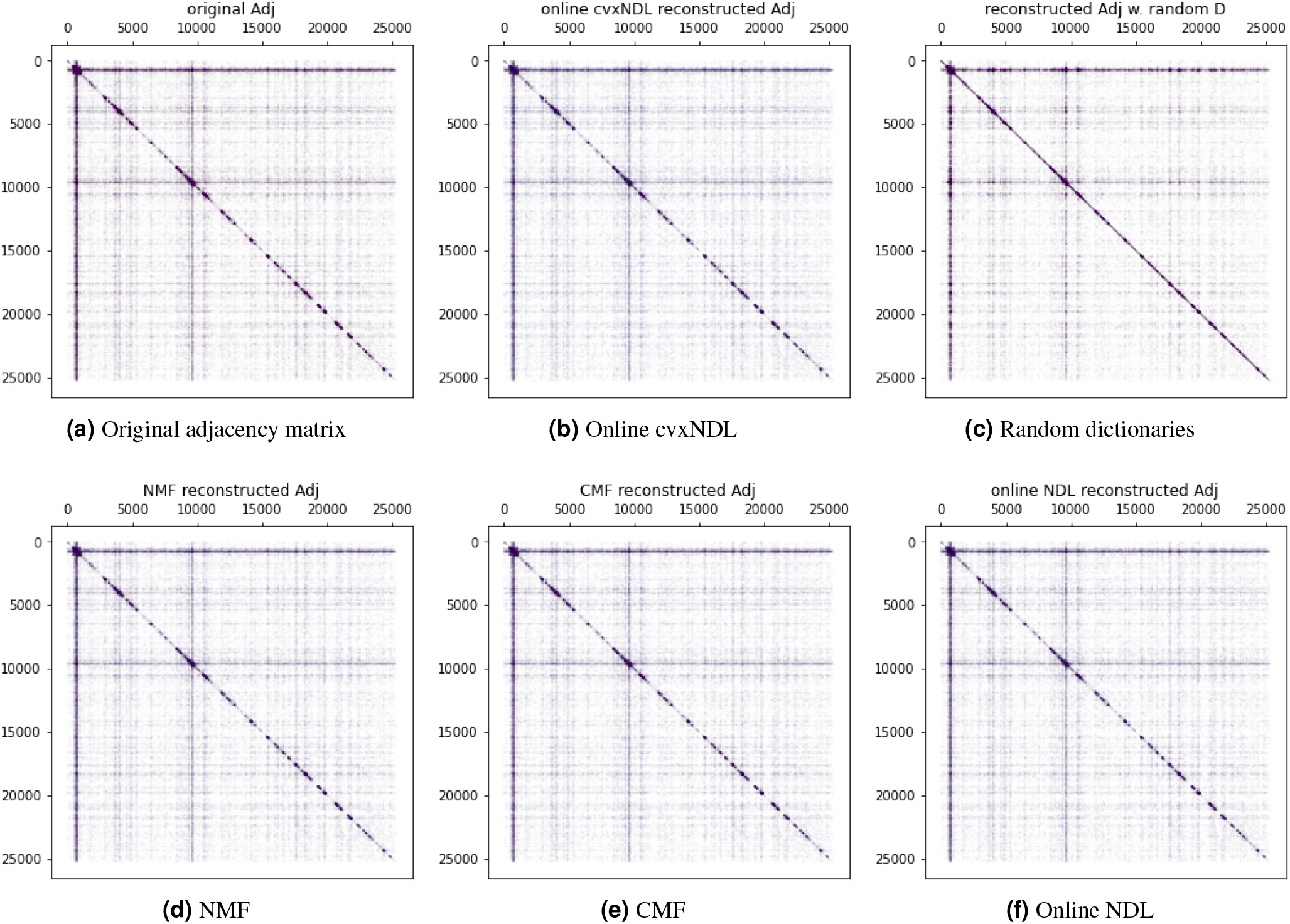
Reconstructed network comparisons based on different baseline methods and random dictionaries, applied on *Drosophila* chromosome 3L ChIA-Drop Data. (a): The original adjacency matrix. (b, c, d, e, f): Reconstructed network adjacency matrices with online cxvNDL, random dictionary elements, NMF, CMF and online NDL.

**Figure 19.**
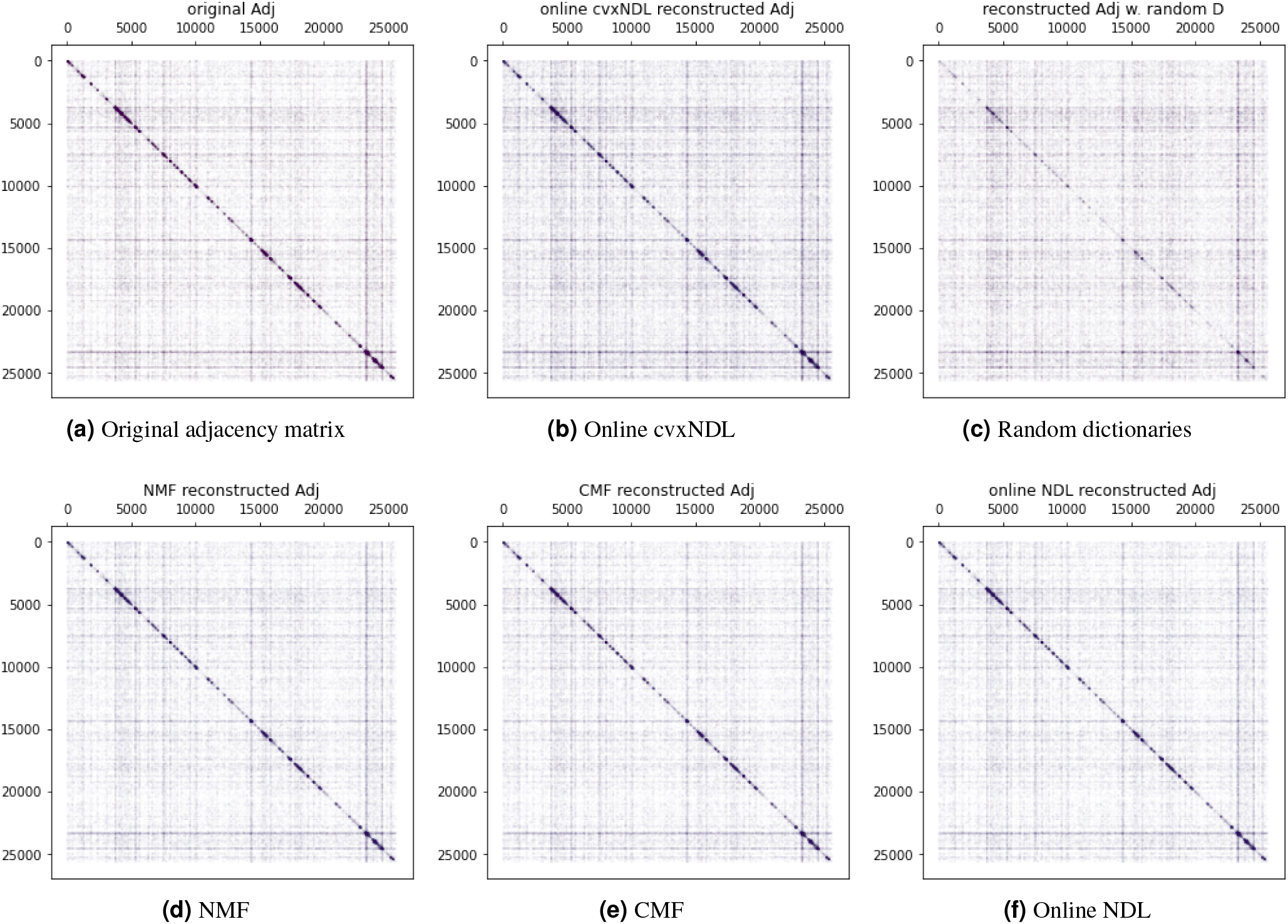
Reconstructed network comparisons based on different baseline methods and random dictionaries, applied on *Drosophila* chromosome 3R ChIA-Drop Data. (a): The original adjacency matrix. (b, c, d, e, f): Reconstructed network adjacency with online cxvNDL, random dictionary elements, NMF, CMF and online NDL.

### 5.5 Gene Ontology Enrichment Analysis

To associate a biological function with each dictionary element, we performed a gene ontology (GO) enrichment analysis for each element and the corresponding chromosome. Recall that as a results of the convexity constraint, every dictionary element has its corresponding set of representatives that capture real observed subgraphs which can be mapped back to actual genomic locations. Of most interest is the set of genes that covers at least one vertex in at least one of the representatives, as described in Figure 20. Using the set of representative genes, we run the GO enrichment analysis in http://geneontology.org under annotation setting”Biological process” and reference list “Drosophila Melanogaster,” for each dictionary element. For further analysis, we only selected results with false discovery rate (FDR) < 0.05 and hence obtained candidate sets of enriched GO terms. Note that there may be inherently enriched GO terms for each dictionary element due to the sampling bias. To remove this bias, we ran another GO enrichment analysis with all genes on each chromosome and used that results to filter out the background GO terms for each dictionary element.

**Figure 20.**
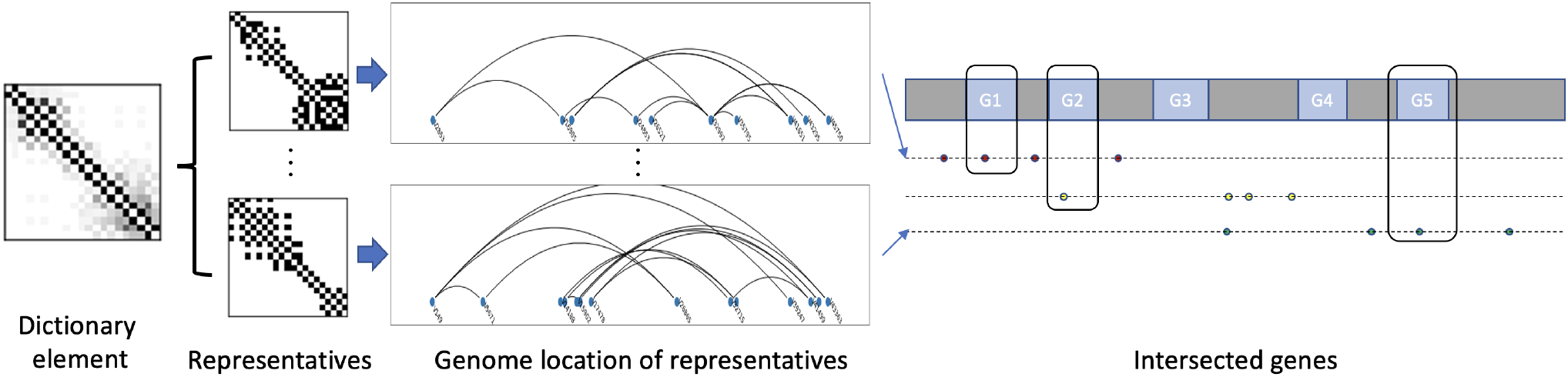
GO enrichment analysis workflow. Each dictionary element is associated with a collection of real subnetwork representatives. These comprise nodes that can be mapped to the genome to identify their locations. A gene is said to cover the node if the DNA fragment corresponding to the node is fully contained within the gene.

We also used the hierarchical structure of GO terms^34^, in which all GO terms are nodes in a directed acyclic graph and edges indicates their relationship. A child GO term is considered more specific than and parent GO term. Since the GO graph is not a strict hierarchy (a child node may have multiple parent nodes), to further improve the results, we performed the following processing. For each GO term: i) we first found all the paths between the term and the root node (which is “Biological process” in our setting), and ii) we removed all intermediate parent GO terms from its enriched GO terms set. By iteratively repeating this filtering process for each dictionary element, we arrived at a set of most specific GO terms for each dictionary element.

#### 5.5.1 Dictionary Elements Associated with GO Terms

We investigated the most frequently enriched GO terms as well as the least frequently enriched GO terms for each chromosome, and identified the corresponding dictionary elements where they were found to be enriched. The results are shown in Tables 5 to 8. For each dictionary element, we computed its density (complexity) *ρ* via 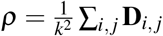 and the median genome distance between all consecutive pairs of nodes, denoted by *d*_med_. The full set of results for the densities and median distances for all dictionary elements and all chromosomes is provided in Tables 15 and 16.

**Table 5.** Top-5 enriched and least enriched GO terms, i.e., terms that most frequently occur as representative in dictionary elements of chr2L. Column ‘#’ indicates the number of dictionary elements that show enrichment of the GO term. We also report the importance scores along with the density of dictionary element *ρ* and median distance of all consecutive pairs of nodes in its representatives *d*_med_.

| most frequent GO term | # | top 3 dictionaries | least frequent GO term | # | dictionary |
| --- | --- | --- | --- | --- | --- |
| (GO:2000241) regulation of reproductive process | 5 | dict_2 (0.085)<br>dict_21 (0.070)<br>dict_6 (0.044)<br>$\rho=0.134, 0.142, 0.161$<br>$d_{\text{med}}=9906, 8105, 10024$ | (GO:0007485) imaginal disc-derived male genitalia development | 1 | dict_21 (0.070)<br>$\rho=0.142$<br>$d_{\text{med}}=8105$ |
| (GO:0046716) muscle cell cellular homeostasis | 4 | dict_14 (0.055)<br>dict_6 (0.044)<br>dict_12 (0.029)<br>$\rho=0.141, 0.161, 0.203$<br>$d_{\text{med}}=10928, 10024, 9979$ | (GO:0008347) glial cell migration | 1 | dict_5 (0.074)<br>$\rho=0.132$<br>$d_{\text{med}}=8547$ |
| (GO:0007422) peripheral nervous system development | 3 | dict_5 (0.074)<br>dict_7 (0.061)<br>dict_8 (0.057)<br>$\rho=0.132, 0.158, 0.147$<br>$d_{\text{med}}=8547, 8870, 10692$ | (GO:0002920) regulation of humoral immune response | 1 | dict_21 (0.070)<br>$\rho=0.142$<br>$d_{\text{med}}=8105$ |
| (GO:0071310) cellular response to organic substance | 3 | dict_2 (0.085)<br>dict_21 (0.070)<br>dict_7 (0.061)<br>$\rho=0.134, 0.142, 0.158$<br>$d_{\text{med}}=9906, 8105, 8870$ | (GO:0016075) rRNA catabolic process | 1 | dict_8 (0.057)<br>$\rho=0.147$<br>$d_{\text{med}}=10692$ |
| (GO:0007400) neuroblast fate determination | 3 | dict_5 (0.074)<br>dict_21 (0.070)<br>dict_8 (0.057)<br>$\rho=0.132, 0.142, 0.147$<br>$d_{\text{med}}=8547, 8105, 10692$ | (GO:0008258) head involution | 1 | dict_8 (0.057)<br>$\rho=0.147$<br>$d_{\text{med}}=10692$ |

**Table 6.**
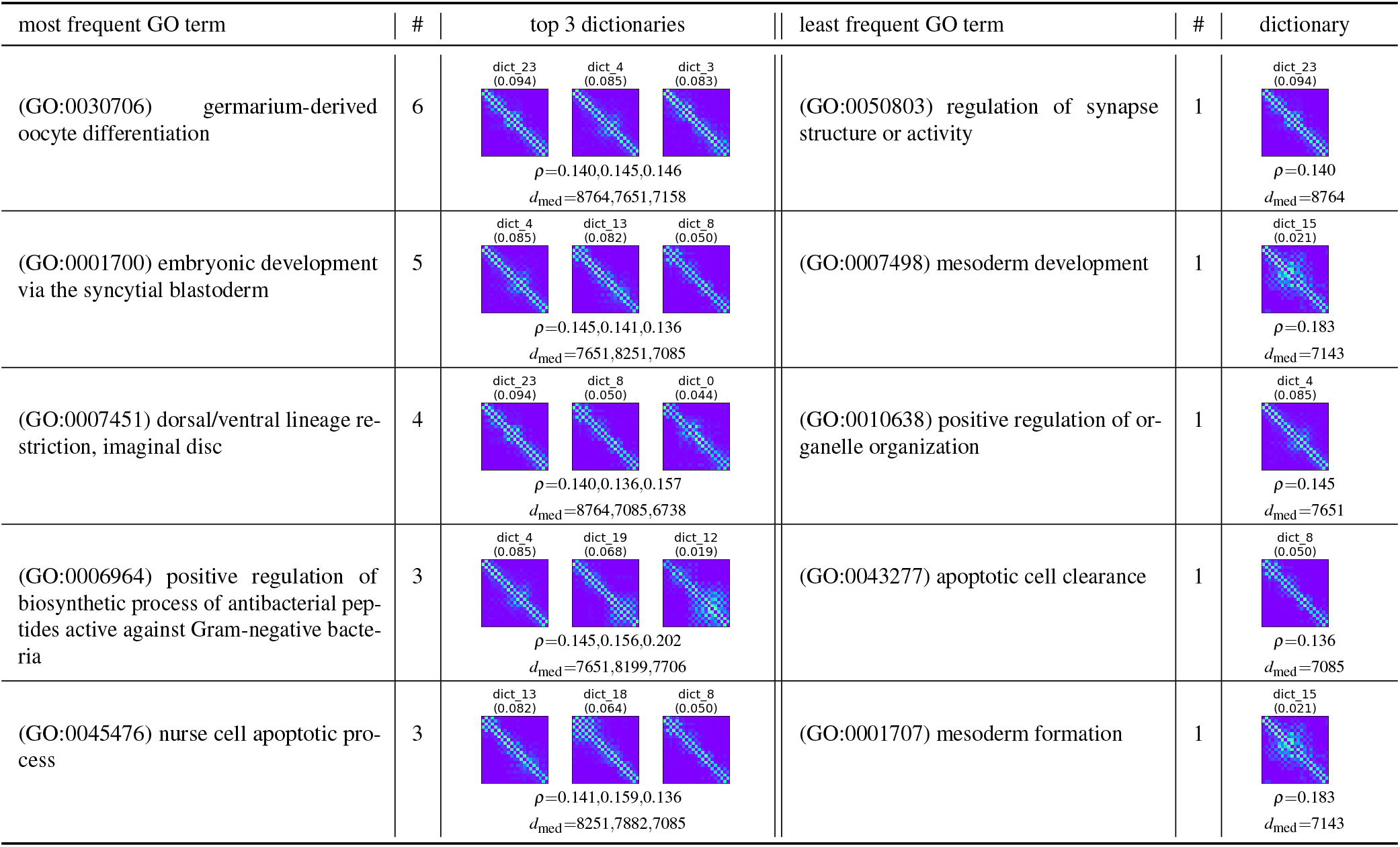
Top-5 enriched and least enriched GO terms, i.e., terms that most frequently occur as representative in dictionary elements of chr2R. Column ‘#’ indicates the number of dictionary elements that show enrichment of the GO term. We also report the importance scores along with the density of dictionary element *ρ* and median distance of all consecutive pairs of nodes in its representatives *d*_med_.

| most frequent GO term | # | top 3 dictionaries | least frequent GO term | # | dictionary |
| --- | --- | --- | --- | --- | --- |
| (GO:0030706) germarium-derived oocyte differentiation | 6 | dict_23 (0.094)<br>dict_4 (0.085)<br>dict_3 (0.083)<br>$\rho=0.140, 0.145, 0.146$<br>$d_{\text{med}}=8764, 7651, 7158$ | (GO:0050803) regulation of synapse structure or activity | 1 | dict_23 (0.094)<br>$\rho=0.140$<br>$d_{\text{med}}=8764$ |
| (GO:0001700) embryonic development via the syncytial blastoderm | 5 | dict_4 (0.085)<br>dict_13 (0.082)<br>dict_8 (0.050)<br>$\rho=0.145, 0.141, 0.136$<br>$d_{\text{med}}=7651, 8251, 7085$ | (GO:0007498) mesoderm development | 1 | dict_15 (0.021)<br>$\rho=0.183$<br>$d_{\text{med}}=7143$ |
| (GO:0007451) dorsal/ventral lineage restriction, imaginal disc | 4 | dict_23 (0.094)<br>dict_8 (0.050)<br>dict_0 (0.044)<br>$\rho=0.140, 0.136, 0.157$<br>$d_{\text{med}}=8764, 7085, 6738$ | (GO:0010638) positive regulation of organelle organization | 1 | dict_4 (0.085)<br>$\rho=0.145$<br>$d_{\text{med}}=7651$ |
| (GO:0006964) positive regulation of biosynthetic process of antibacterial peptides active against Gram-negative bacteria | 3 | dict_4 (0.085)<br>dict_19 (0.068)<br>dict_12 (0.019)<br>$\rho=0.145, 0.156, 0.202$<br>$d_{\text{med}}=7651, 8199, 7706$ | (GO:0043277) apoptotic cell clearance | 1 | dict_8 (0.050)<br>$\rho=0.136$<br>$d_{\text{med}}=7085$ |
| (GO:0045476) nurse cell apoptotic process | 3 | dict_13 (0.082)<br>dict_18 (0.064)<br>dict_8 (0.050)<br>$\rho=0.141, 0.159, 0.136$<br>$d_{\text{med}}=8251, 7882, 7085$ | (GO:0001707) mesoderm formation | 1 | dict_15 (0.021)<br>$\rho=0.183$<br>$d_{\text{med}}=7143$ |

**Table 7.**
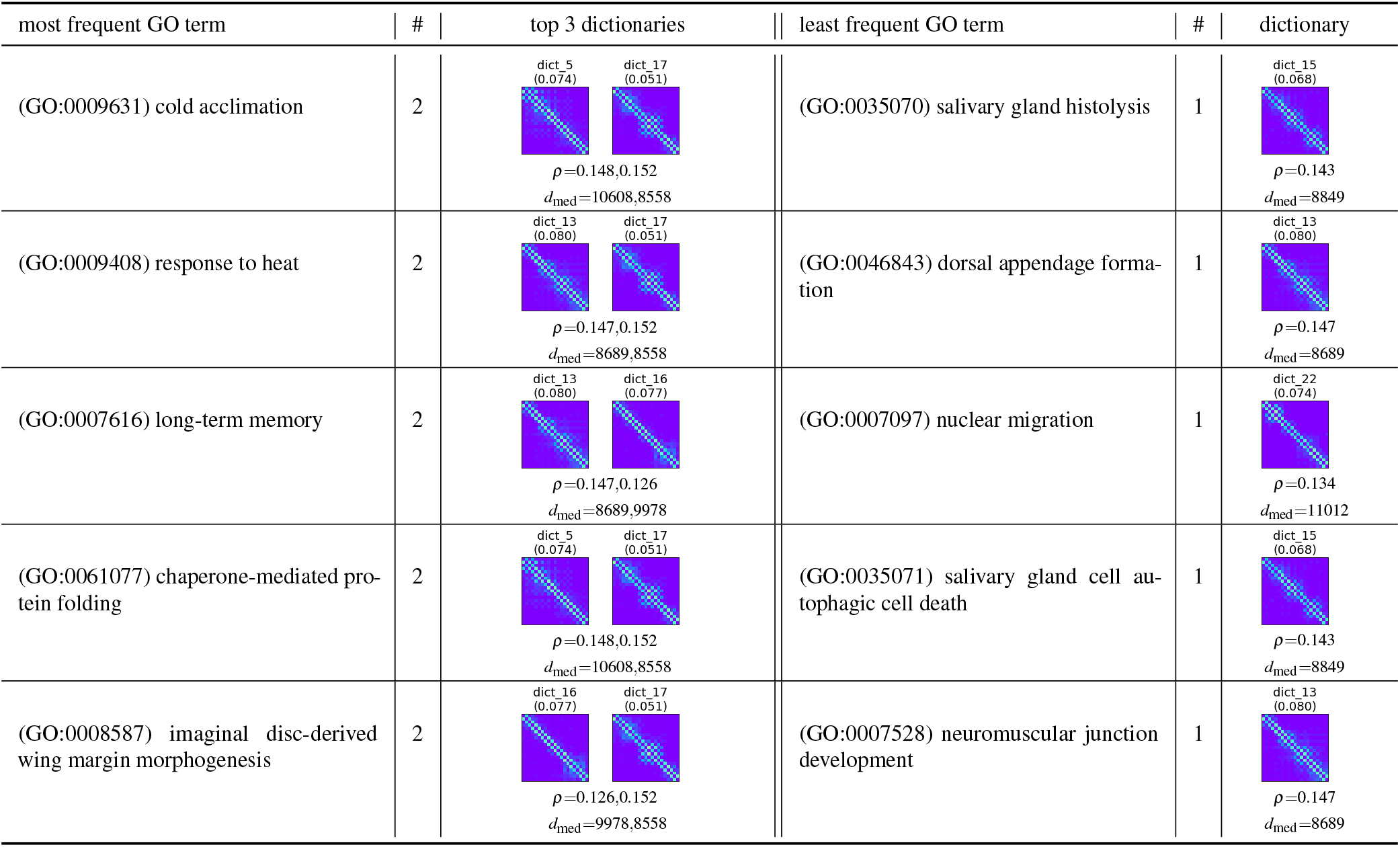
Top-5 enriched and least enriched GO terms, i.e., terms that most frequently occur as representative in dictionary elements of chr3L. Column ‘#’ indicates the number of dictionary elements that show enrichment of the GO term. We also report the importance scores along with the density of dictionary element *ρ* and median distance of all consecutive pairs of nodes in its representatives *d*_med_.

| most frequent GO term | # | top 3 dictionaries | least frequent GO term | # | dictionary |
| --- | --- | --- | --- | --- | --- |
| (GO:0009631) cold acclimation | 2 | dict_5<br>(0.074)<br>dict_17<br>(0.051)<br>$\rho=0.148, 0.152$<br>$d_{med}=10608, 8558$ | (GO:0035070) salivary gland histolysis | 1 | dict_15<br>(0.068)<br>$\rho=0.143$<br>$d_{med}=8849$ |
| (GO:0009408) response to heat | 2 | dict_13<br>(0.080)<br>dict_17<br>(0.051)<br>$\rho=0.147, 0.152$<br>$d_{med}=8689, 8558$ | (GO:0046843) dorsal appendage formation | 1 | dict_13<br>(0.080)<br>$\rho=0.147$<br>$d_{med}=8689$ |
| (GO:0007616) long-term memory | 2 | dict_13<br>(0.080)<br>dict_16<br>(0.077)<br>$\rho=0.147, 0.126$<br>$d_{med}=8689, 9978$ | (GO:0007097) nuclear migration | 1 | dict_22<br>(0.074)<br>$\rho=0.134$<br>$d_{med}=11012$ |
| (GO:0061077) chaperone-mediated protein folding | 2 | dict_5<br>(0.074)<br>dict_17<br>(0.051)<br>$\rho=0.148, 0.152$<br>$d_{med}=10608, 8558$ | (GO:0035071) salivary gland cell autophagic cell death | 1 | dict_15<br>(0.068)<br>$\rho=0.143$<br>$d_{med}=8849$ |
| (GO:0008587) imaginal disc-derived wing margin morphogenesis | 2 | dict_16<br>(0.077)<br>dict_17<br>(0.051)<br>$\rho=0.126, 0.152$<br>$d_{med}=9978, 8558$ | (GO:0007528) neuromuscular junction development | 1 | dict_13<br>(0.080)<br>$\rho=0.147$<br>$d_{med}=8689$ |

**Table 8.** Top-5 enriched and least enriched GO terms, i.e., terms that most frequently occur as representative in dictionary elements of chr3R. Column ‘#’ indicates the number of dictionary elements that show enrichment of the GO term. We also report the importance scores along with the density of dictionary element *ρ* and median distance of all consecutive pairs of nodes in its representatives *d*_med_.

| most frequent GO term | # | top 3 dictionaries | least frequent GO term | # | dictionary |
| --- | --- | --- | --- | --- | --- |
| (GO:0001819) positive regulation of cytokine production | 7 | dict_20 (0.121)<br>dict_7 (0.059)<br>dict_9 (0.049)<br>$\rho=0.126, 0.146, 0.157$<br>$d_{med}=12791, 12830, 11930$ | (GO:0061448) connective tissue development | 1 | dict_12 (0.085)<br>$\rho=0.142$<br>$d_{med}=13455$ |
| (GO:0008015) blood circulation | 7 | dict_20 (0.121)<br>dict_12 (0.085)<br>dict_4 (0.066)<br>$\rho=0.126, 0.142, 0.138$<br>$d_{med}=12791, 13455, 13674$ | (GO:0051282) regulation of sequestering of calcium ion | 1 | dict_20 (0.121)<br>$\rho=0.126$<br>$d_{med}=12791$ |
| (GO:0045948) positive regulation of translational initiation | 5 | dict_20 (0.121)<br>dict_4 (0.066)<br>dict_14 (0.049)<br>$\rho=0.126, 0.138, 0.162$<br>$d_{med}=12791, 13674, 12572$ | (GO:0043123) positive regulation of I-kappaB kinase/NF-kappaB signaling | 1 | dict_13 (0.016)<br>$\rho=0.204$<br>$d_{med}=12540$ |
| (GO:0042177) negative regulation of protein catabolic process | 5 | dict_20 (0.121)<br>dict_12 (0.085)<br>dict_4 (0.066)<br>$\rho=0.126, 0.142, 0.138$<br>$d_{med}=12791, 13455, 13674$ | (GO:0007435) salivary gland morphogenesis | 1 | dict_13 (0.016)<br>$\rho=0.204$<br>$d_{med}=12540$ |
| (GO:0043065) positive regulation of apoptotic process | 4 | dict_20 (0.121)<br>dict_7 (0.059)<br>dict_3 (0.041)<br>$\rho=0.126, 0.146, 0.179$<br>$d_{med}=12791, 12830, 11748$ | (GO:0045738) negative regulation of DNA repair | 1 | dict_8 (0.046)<br>$\rho=0.183$<br>$d_{med}=12493$ |

Note that the Drosophila S2 cells are embryonic cells, and most GO terms found are related to cellular reproductive process or developmental process, as expected. From the tables one can also see that different dictionary elements reflect different biological processes and for the same GO term, the dictionary elements share similar patterns. For example, in Table 5, we can see that dictionary elements 19 and 12 share very similar structural patterns, and both of them are enriched in biosynthetic processes of antibacterial peptides. On the other hand, dictionary elements 13 and 8 have a pattern that differs from that of 19 and 12, and they are enriched in dorsal/ventral lineage restriction processes. We also found that dictionary elements with GO term peripheral nervous system development, celluar response to organic substance, and neuroblast fate determination have relatively lower density and smaller median node distances than the top-2 enriched GO terms, *regulation of reproductive process* and *muscle cell cellular homeostasis.* The difference in density and median distance is also reflected by the significantly different dictionary patterns observed, such as for example dictionary element 12 and dictionary element 5; the former element has a much higher density and median distance than the latter.

There are also a few shared GO terms that are enriched in both chr2L and chr2R (11 shared terms in total), and in both chr3L and chr3R (3 shared terms in total). The results are reported in Table 9 and 10. We found that there are very few shared terms between the two chromosomes, when compared to the roughly one hundred uniquely enriched GO terms for each chromosome. Most of the shared terms also have “similar” patterns (which can be seen visually or through a simple computation of the *ℓ*_2_ distance between their flattened adjacency matrices) of their corresponding dictionary elements.

**Table 9.** GO terms shared between chr2L and chr2R.

| GO_term | chr2L dictionaries | chr2R dictionaries |
| --- | --- | --- |
| (GO:0016325) oocyte microtubule cytoskeleton organization | dict_5 (0.074)<br>dict_7 (0.061)<br>dict_6 (0.044) | dict_14 (0.013) |
| (GO:1901701) cellular response to oxygen-containing compound | dict_2 (0.085)<br>dict_7 (0.061) | dict_8 (0.050) |
| (GO:0007298) border follicle cell migration | dict_2 (0.085)<br>dict_21 (0.070) | dict_4 (0.085)<br>dict_3 (0.083)<br>dict_18 (0.064) |
| (GO:0043410) positive regulation of MAPK cascade | dict_2 (0.085)<br>dict_8 (0.057) | dict_4 (0.085)<br>dict_8 (0.050) |
| (GO:0016049) cell growth | dict_21 (0.070) | dict_8 (0.050) |
| (GO:0035331) negative regulation of hippo signaling | dict_8 (0.057) | dict_4 (0.085) |
| (GO:0051962) positive regulation of nervous system development | dict_7 (0.061) | dict_15 (0.021) |
| (GO:0060322) head development | dict_8 (0.057) | dict_4 (0.085) |
| (GO:0007293) germarium-derived egg chamber formation | dict_8 (0.057) | dict_23 (0.094)<br>dict_4 (0.085)<br>dict_13 (0.082)<br>dict_15 (0.021) |
| (GO:0002164) larval development | dict_6 (0.044) | dict_15 (0.021) |
| (GO:0007420) brain development | dict_6 (0.044) | dict_4 (0.085)<br>dict_18 (0.064) |

**Table 10.** GO terms shared between chr3L and chr3R.

| GO_term | chr3L dictionaries | chr3R dictionaries |
| --- | --- | --- |
| (GO:0070373) negative regulation of ERK1 and ERK2 cascade | dict_13 (0.080)<br>dict_22 (0.074)<br>dict_3 (0.045)<br>dict_1 (0.035) | dict_8 (0.046) |
| (GO:0007140) male meiotic nuclear division | dict_23 (0.029) | dict_24 (0.017) |
| (GO:0046777) protein autophosphorylation | dict_22 (0.074) | dict_8 (0.046) |

#### 5.5.2 Additional Results

Here we report more detailed results for each dictionary element, including its number of enriched GO terms (Tables 11, 12, 13, 14), density (Table 15) and median distance (Table 16).

**Table 11.**
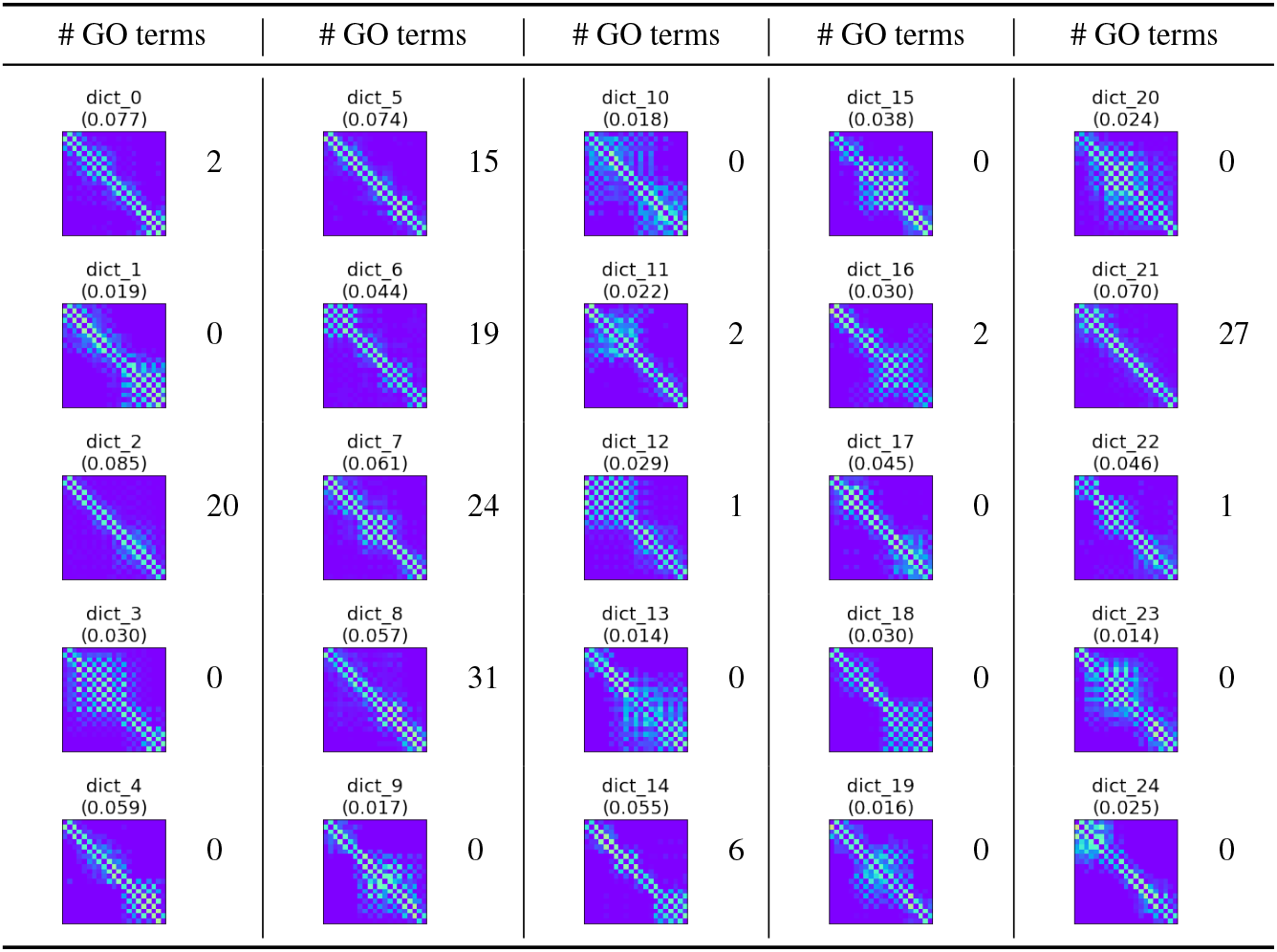
Number of enriched GO terms for each dictionary element identified for chr2L.

| # GO terms | # GO terms | # GO terms | # GO terms | # GO terms |
| --- | --- | --- | --- | --- |
| dict_0<br>(0.077)<br>2 | dict_5<br>(0.074)<br>15 | dict_10<br>(0.018)<br>0 | dict_15<br>(0.038)<br>0 | dict_20<br>(0.024)<br>0 |
| dict_1<br>(0.019)<br>0 | dict_6<br>(0.044)<br>19 | dict_11<br>(0.022)<br>2 | dict_16<br>(0.030)<br>2 | dict_21<br>(0.070)<br>27 |
| dict_2<br>(0.085)<br>20 | dict_7<br>(0.061)<br>24 | dict_12<br>(0.029)<br>1 | dict_17<br>(0.045)<br>0 | dict_22<br>(0.046)<br>1 |
| dict_3<br>(0.030)<br>0 | dict_8<br>(0.057)<br>31 | dict_13<br>(0.014)<br>0 | dict_18<br>(0.030)<br>0 | dict_23<br>(0.014)<br>0 |
| dict_4<br>(0.059)<br>0 | dict_9<br>(0.017)<br>0 | dict_14<br>(0.055)<br>6 | dict_19<br>(0.016)<br>0 | dict_24<br>(0.025)<br>0 |

**Table 12.**
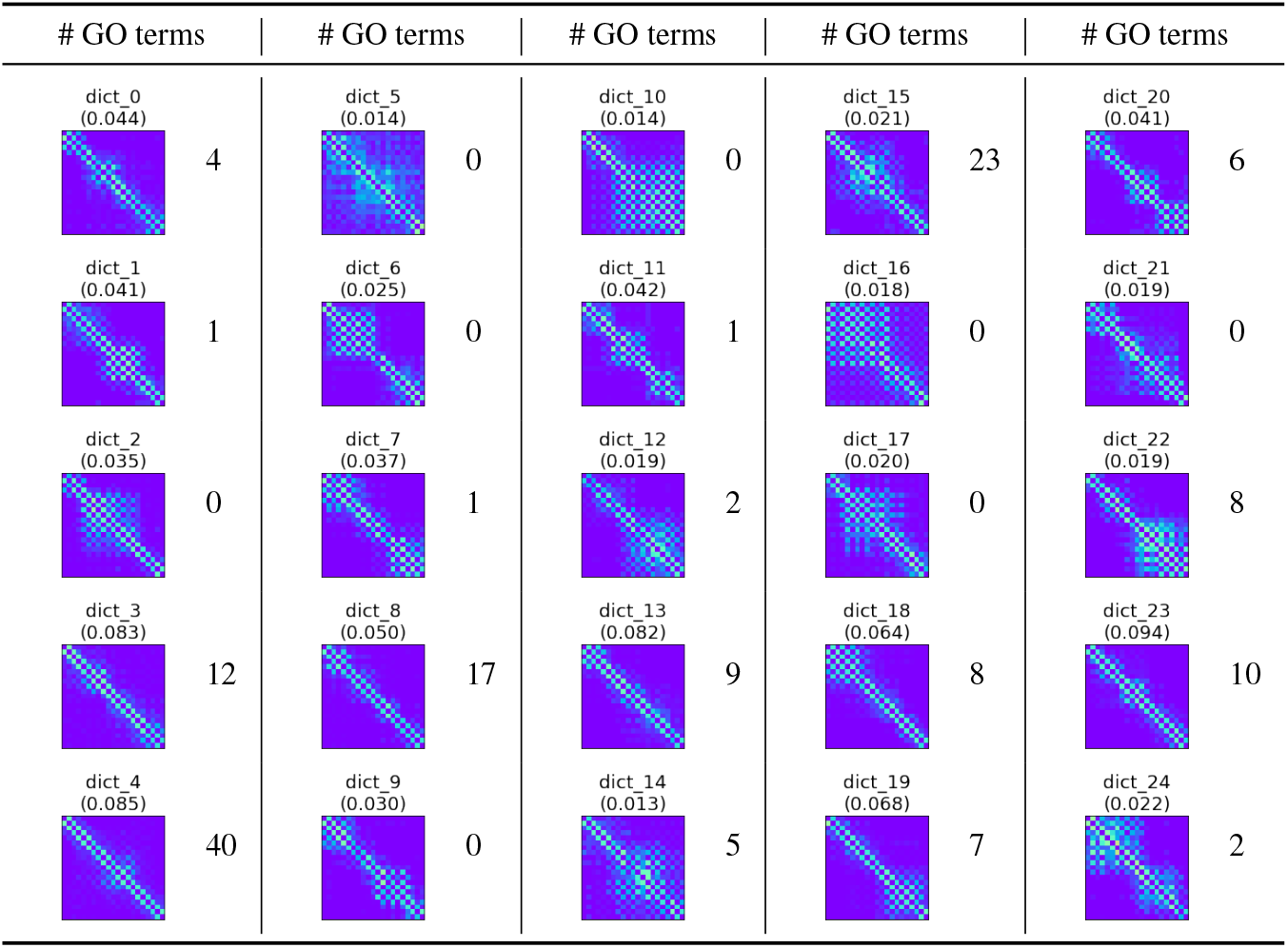
Number of enriched GO terms for each dictionary element identified for chr2R.

| # GO terms | # GO terms | # GO terms | # GO terms | # GO terms |
| --- | --- | --- | --- | --- |
| dict_0<br>(0.044)<br>4 | dict_5<br>(0.014)<br>0 | dict_10<br>(0.014)<br>0 | dict_15<br>(0.021)<br>23 | dict_20<br>(0.041)<br>6 |
| dict_1<br>(0.041)<br>1 | dict_6<br>(0.025)<br>0 | dict_11<br>(0.042)<br>1 | dict_16<br>(0.018)<br>0 | dict_21<br>(0.019)<br>0 |
| dict_2<br>(0.035)<br>0 | dict_7<br>(0.037)<br>1 | dict_12<br>(0.019)<br>2 | dict_17<br>(0.020)<br>0 | dict_22<br>(0.019)<br>8 |
| dict_3<br>(0.083)<br>12 | dict_8<br>(0.050)<br>17 | dict_13<br>(0.082)<br>9 | dict_18<br>(0.064)<br>8 | dict_23<br>(0.094)<br>10 |
| dict_4<br>(0.085)<br>40 | dict_9<br>(0.030)<br>0 | dict_14<br>(0.013)<br>5 | dict_19<br>(0.068)<br>7 | dict_24<br>(0.022)<br>2 |

**Table 13.**
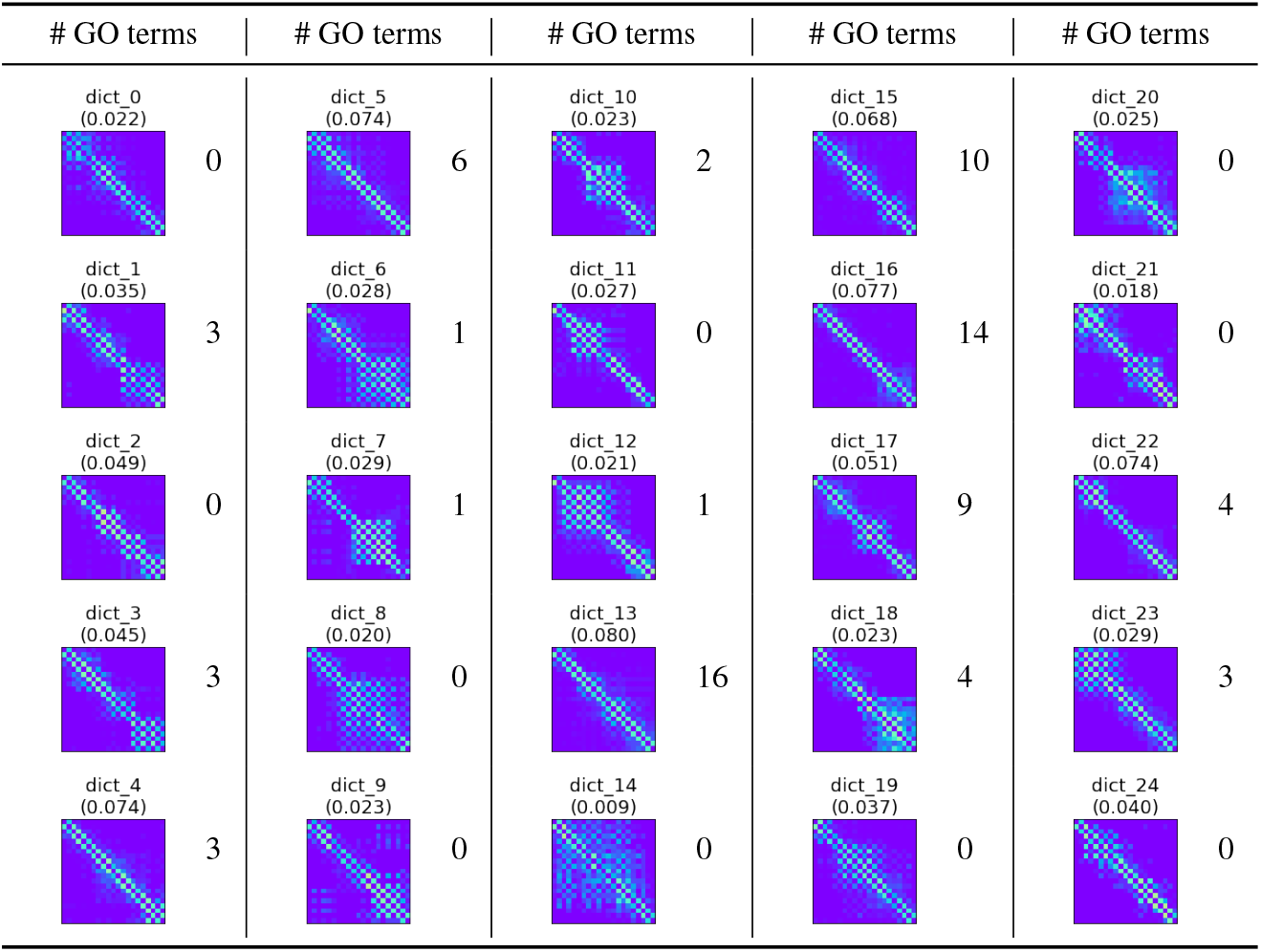
Number of enriched GO terms for each dictionary element identified for chr3L.

| # GO terms | # GO terms | # GO terms | # GO terms | # GO terms |
| --- | --- | --- | --- | --- |
| dict_0<br>(0.022) | dict_5<br>(0.074) | dict_10<br>(0.023) | dict_15<br>(0.068) | dict_20<br>(0.025) |
| 0 | 6 | 2 | 10 | 0 |
| dict_1<br>(0.035) | dict_6<br>(0.028) | dict_11<br>(0.027) | dict_16<br>(0.077) | dict_21<br>(0.018) |
| 3 | 1 | 0 | 14 | 0 |
| dict_2<br>(0.049) | dict_7<br>(0.029) | dict_12<br>(0.021) | dict_17<br>(0.051) | dict_22<br>(0.074) |
| 0 | 1 | 1 | 9 | 4 |
| dict_3<br>(0.045) | dict_8<br>(0.020) | dict_13<br>(0.080) | dict_18<br>(0.023) | dict_23<br>(0.029) |
| 3 | 0 | 16 | 4 | 3 |
| dict_4<br>(0.074) | dict_9<br>(0.023) | dict_14<br>(0.009) | dict_19<br>(0.037) | dict_24<br>(0.040) |
| 3 | 0 | 0 | 0 | 0 |

**Table 14.**
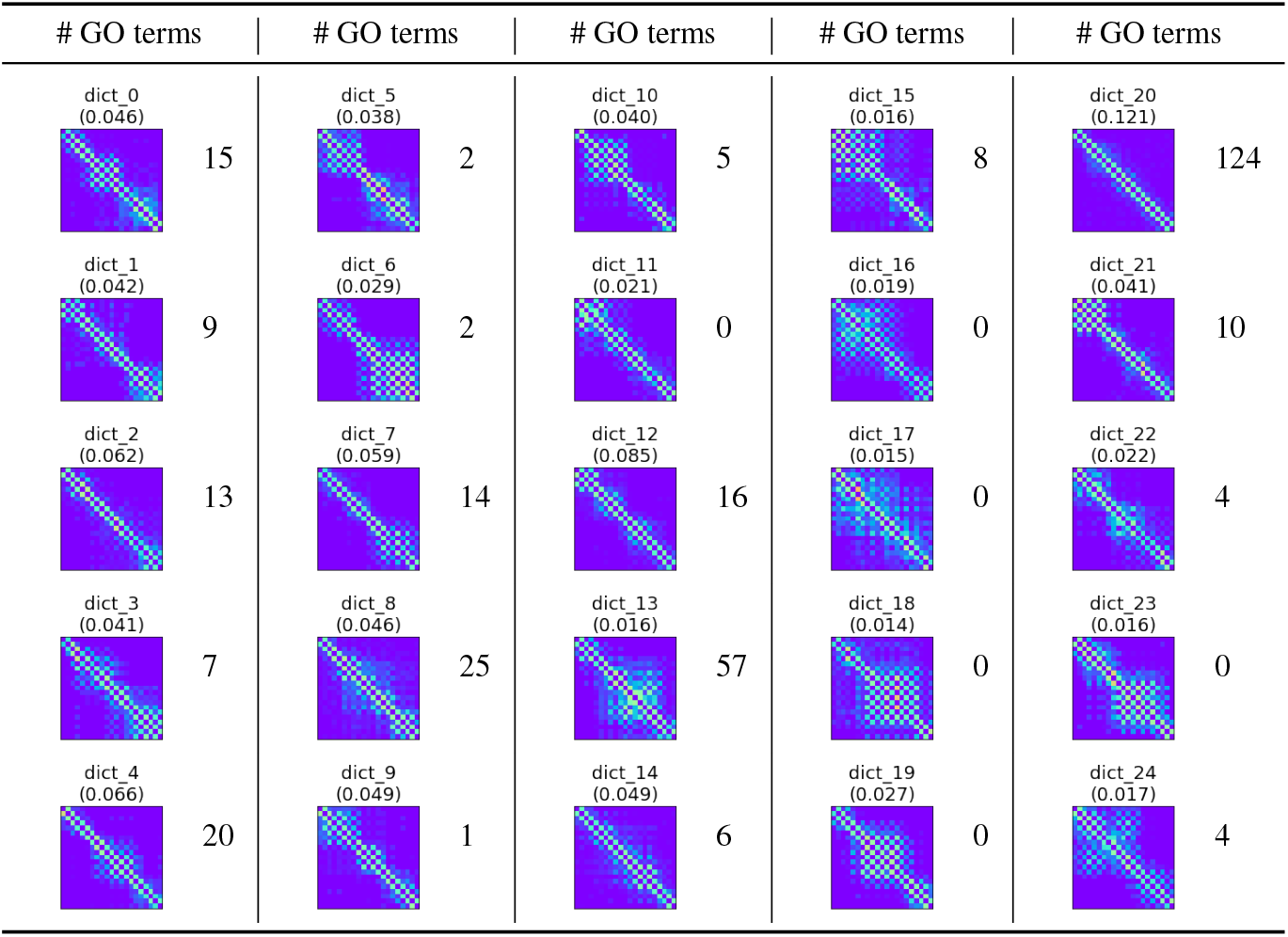
Number of enriched GO terms for each dictionary element identified for chr3R.

| # GO terms | # GO terms | # GO terms | # GO terms | # GO terms |
| --- | --- | --- | --- | --- |
| dict_0<br>(0.046) | dict_5<br>(0.038) | dict_10<br>(0.040) | dict_15<br>(0.016) | dict_20<br>(0.121) |
| 15 | 2 | 5 | 8 | 124 |
| dict_1<br>(0.042) | dict_6<br>(0.029) | dict_11<br>(0.021) | dict_16<br>(0.019) | dict_21<br>(0.041) |
| 9 | 2 | 0 | 0 | 10 |
| dict_2<br>(0.062) | dict_7<br>(0.059) | dict_12<br>(0.085) | dict_17<br>(0.015) | dict_22<br>(0.022) |
| 13 | 14 | 16 | 0 | 4 |
| dict_3<br>(0.041) | dict_8<br>(0.046) | dict_13<br>(0.016) | dict_18<br>(0.014) | dict_23<br>(0.016) |
| 7 | 25 | 57 | 0 | 0 |
| dict_4<br>(0.066) | dict_9<br>(0.049) | dict_14<br>(0.049) | dict_19<br>(0.027) | dict_24<br>(0.017) |
| 20 | 1 | 6 | 0 | 4 |

**Table 15.** Density of dictionary elements, reported for all chromosomes.

| Dictionary element | chr2L | chr2R | chr3L | chr3R |
| --- | --- | --- | --- | --- |
| 1 | 0.146 | 0.158 | 0.168 | 0.161 |
| 2 | 0.188 | 0.165 | 0.156 | 0.157 |
| 3 | 0.134 | 0.185 | 0.141 | 0.140 |
| 4 | 0.220 | 0.147 | 0.159 | 0.179 |
| 5 | 0.145 | 0.146 | 0.142 | 0.139 |
| 6 | 0.132 | 0.297 | 0.148 | 0.173 |
| 7 | 0.162 | 0.189 | 0.191 | 0.184 |
| 8 | 0.158 | 0.184 | 0.164 | 0.147 |
| 9 | 0.148 | 0.136 | 0.210 | 0.183 |
| 10 | 0.177 | 0.166 | 0.168 | 0.157 |
| 11 | 0.220 | 0.261 | 0.163 | 0.161 |
| 12 | 0.168 | 0.162 | 0.145 | 0.157 |
| 13 | 0.204 | 0.203 | 0.186 | 0.142 |
| 14 | 0.225 | 0.142 | 0.148 | 0.205 |
| 15 | 0.142 | 0.229 | 0.262 | 0.163 |
| 16 | 0.173 | 0.184 | 0.143 | 0.205 |
| 17 | 0.189 | 0.263 | 0.127 | 0.224 |
| 18 | 0.161 | 0.219 | 0.152 | 0.251 |
| 19 | 0.182 | 0.159 | 0.183 | 0.242 |
| 20 | 0.187 | 0.156 | 0.170 | 0.193 |
| 21 | 0.231 | 0.157 | 0.199 | 0.126 |
| 22 | 0.143 | 0.195 | 0.165 | 0.150 |
| 23 | 0.162 | 0.201 | 0.134 | 0.175 |
| 24 | 0.223 | 0.141 | 0.167 | 0.212 |
| 25 | 0.167 | 0.212 | 0.140 | 0.208 |

**Table 16.**
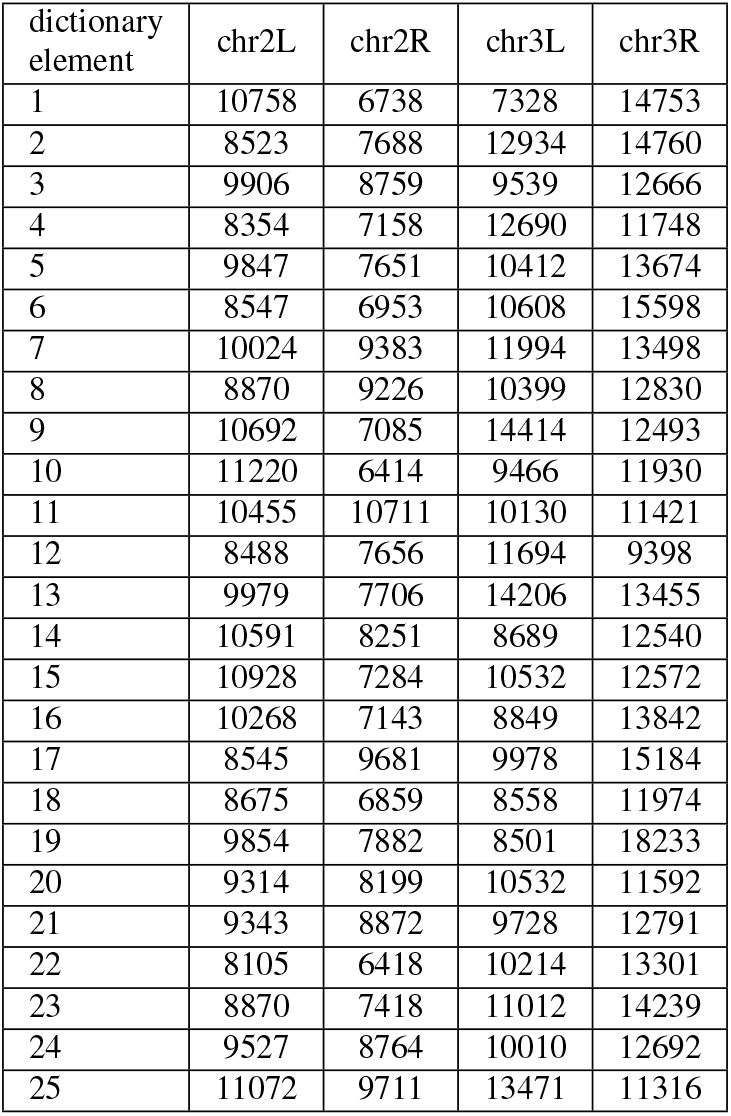
Median distance of pairwise interacting nodes within each dictionary element and for each chromosome.

| dictionary element | chr2L | chr2R | chr3L | chr3R |
| --- | --- | --- | --- | --- |
| 1 | 10758 | 6738 | 7328 | 14753 |
| 2 | 8523 | 7688 | 12934 | 14760 |
| 3 | 9906 | 8759 | 9539 | 12666 |
| 4 | 8354 | 7158 | 12690 | 11748 |
| 5 | 9847 | 7651 | 10412 | 13674 |
| 6 | 8547 | 6953 | 10608 | 15598 |
| 7 | 10024 | 9383 | 11994 | 13498 |
| 8 | 8870 | 9226 | 10399 | 12830 |
| 9 | 10692 | 7085 | 14414 | 12493 |
| 10 | 11220 | 6414 | 9466 | 11930 |
| 11 | 10455 | 10711 | 10130 | 11421 |
| 12 | 8488 | 7656 | 11694 | 9398 |
| 13 | 9979 | 7706 | 14206 | 13455 |
| 14 | 10591 | 8251 | 8689 | 12540 |
| 15 | 10928 | 7284 | 10532 | 12572 |
| 16 | 10268 | 7143 | 8849 | 13842 |
| 17 | 8545 | 9681 | 9978 | 15184 |
| 18 | 8675 | 6859 | 8558 | 11974 |
| 19 | 9854 | 7882 | 8501 | 18233 |
| 20 | 9314 | 8199 | 10532 | 11592 |
| 21 | 9343 | 8872 | 9728 | 12791 |
| 22 | 8105 | 6418 | 10214 | 13301 |
| 23 | 8870 | 7418 | 11012 | 14239 |
| 24 | 9527 | 8764 | 10010 | 12692 |
| 25 | 11072 | 9711 | 13471 | 11316 |

